# *Pseudomonas* effector AvrC is a rhamnosyltransferase with broad substrate specificity

**DOI:** 10.64898/2026.08.26.747443

**Authors:** Nalleli Payne, Kelly A. Servage, Kim Orth, Jessie Fernandez, Wei Peng

**Author notes:** Corresponding author: Wei Peng. These authors contributed equally to this work (Nalleli Payne, Wei Peng).

## Abstract

Plant cells can directly or indirectly detect bacterial effectors, triggering a hypersensitive response to defend against pathogen infection. One extensively studied effector, AvrB, is a Fido (<u>Fi</u>c, <u>Do</u>c, AvrB) domain-containing protein that acts as a glycosyltransferase. AvrC is an elusive *Pseudomonas syringae* avirulence effector protein with significant sequence and structural similarity to AvrB. Combining biochemistry, mass spectrometry, and AlphaFold prediction tools, we show that AvrC is a glycosyltransferase with auto-rhamnosylation activity. Like AvrB, AvrC can rhamnosylate a threonine residue (T166) on the *A. thaliana* “guardee” protein RIN4. *In vitro* assays revealed rhamnosylation substrates for AvrC also include plant coatomer subunits COPE1 and COPZ1. Collectively, our findings indicate that AvrC is a rhamnosyltransferase with broad substrate specificity. Our experimental strategies and findings provide valuable insights into future studies on the characterization of other Fido proteins.

## Introduction

Plant cells use effector-triggered immunity called the hypersensitive response to defend against bacterial infection (1–4). In this mechanism, an avirulent effector protein is recognized by a specific resistance protein in a gene-for-gene manner to elicit programmed cell death (1–4). AvrB is a prototypical effector from the plant pathogen *P. syringae* and belongs to the Fido (<u>Fi</u>c, <u>Do</u>c, and AvrB) protein family (5–11). Recently, we characterized AvrB as a glycosyltransferase (11) and established AvrB as the founding member of Glycosyltransferase Family 138 (GT138) (12, 13). Using UDP-rhamnose as a co-substrate, AvrB rhamnosylates a threonine residue (T166) of the *A. thaliana* “guardee” protein RIN4 (RPM1-interacting protein 4) (11). Modification of RIN4 by AvrB leads to activation of the resistance protein RPM1 (resistance to *Pseudomonas syringae* pv. *maculicola* 1), which subsequently assembles into an oligomeric resistosome complex (2, 3, 9).

AvrC is another *P. syringae* avirulence effector protein that shows significant sequence and structural similarity to AvrB, but AvrC induces different disease phenotypes on soybean cultivars (5, 14, 15). However, limited studies have been performed on AvrC to demonstrate how the effector manipulates host cell signaling at the molecular level. Using a variety of biochemistry, mass spectrometry, and AlphaFold prediction tools, we have characterized the molecular and biochemical functions of AvrC. Our findings reveal that AvrC is a glycosyltransferase that has rhamnosylation activity with broad substrate specificity. AvrC auto-rhamnosylates at a serine residue (S14) when expressed in the *E. coli* strain BL21 (DE3). AvrC also binds to the *A. thaliana* “guardee” protein RIN4 and rhamnosylates T166. Importantly, AvrC can modify other host proteins, such as coatomer complex subunits COPE1 and COPZ1. Our observations provide insights into host cell manipulation by AvrC. The adopted experimental strategies shed light on future investigations for other Fido enzymes.

## Results

### AvrC is a glycosyltransferase that auto-rhamnosylates at S14

AvrC is a bacterial effector containing 352 amino acids with ∼46% of sequence identity compared to AvrB (Fig. S1A, S1B) (14). The predicted AlphaFold structure model confidently shows AvrC adopts a Fido fold that aligns well with AvrB (Fig. S1C, S1D). Hence, we hypothesized that AvrC may be a rhamnosyltransferase like AvrB. Indeed, previously we observed that recombinant wild-type AvrC was modified with a mass shift of +146 Da, consistent with auto-rhamnosylation (11). As expected, mutation of the predicted catalytic arginine 297 (corresponding to R266 of AvrB, Fig. S1A) to an alanine abrogated the auto-catalytic activity of AvrC (11).

To identify the auto-rhamnosylation site on AvrC, we performed a post-translational modification (PTM) search using liquid chromatography-tandem mass spectrometry (LC-MS/MS) after digesting the wild-type AvrC protein with various proteases. Multiple peptides were found to carry the +146 Da modification (Fig. 1A). Among these peptides, we noticed an overlapping eight amino acid region (S11-F17) to be the most significant modification site with the highest numbers of positive peptide hits. We therefore tested a N-terminal deletion construct (or the core Fido domain, AvrC^Δ1-51^) and found that this truncated construct lost auto-modification as expected (Fig. 1B). The AlphaFold structure model of AvrC does not confidently model the position of the N-terminus with respect to the core Fido domain (Fig. S1C, S1D). However, prediction of the N-terminal peptide, AvrC^1-51^, in complex with the core Fido domain AvrC^Δ1-51^ positions S14 in the catalytic site of AvrC (Fig. 1C-1F). We propose that S14 likely receives rhamnose from UDP-rhamnose, as indicated by structure alignment of the AvrC^1-51^-AvrC^Δ1-51^ AlphaFold model against the experimental AvrB complex with UDP-rhamnose and RIN4 (Fig. 1E, 1F) (11).

**Figure 1.**
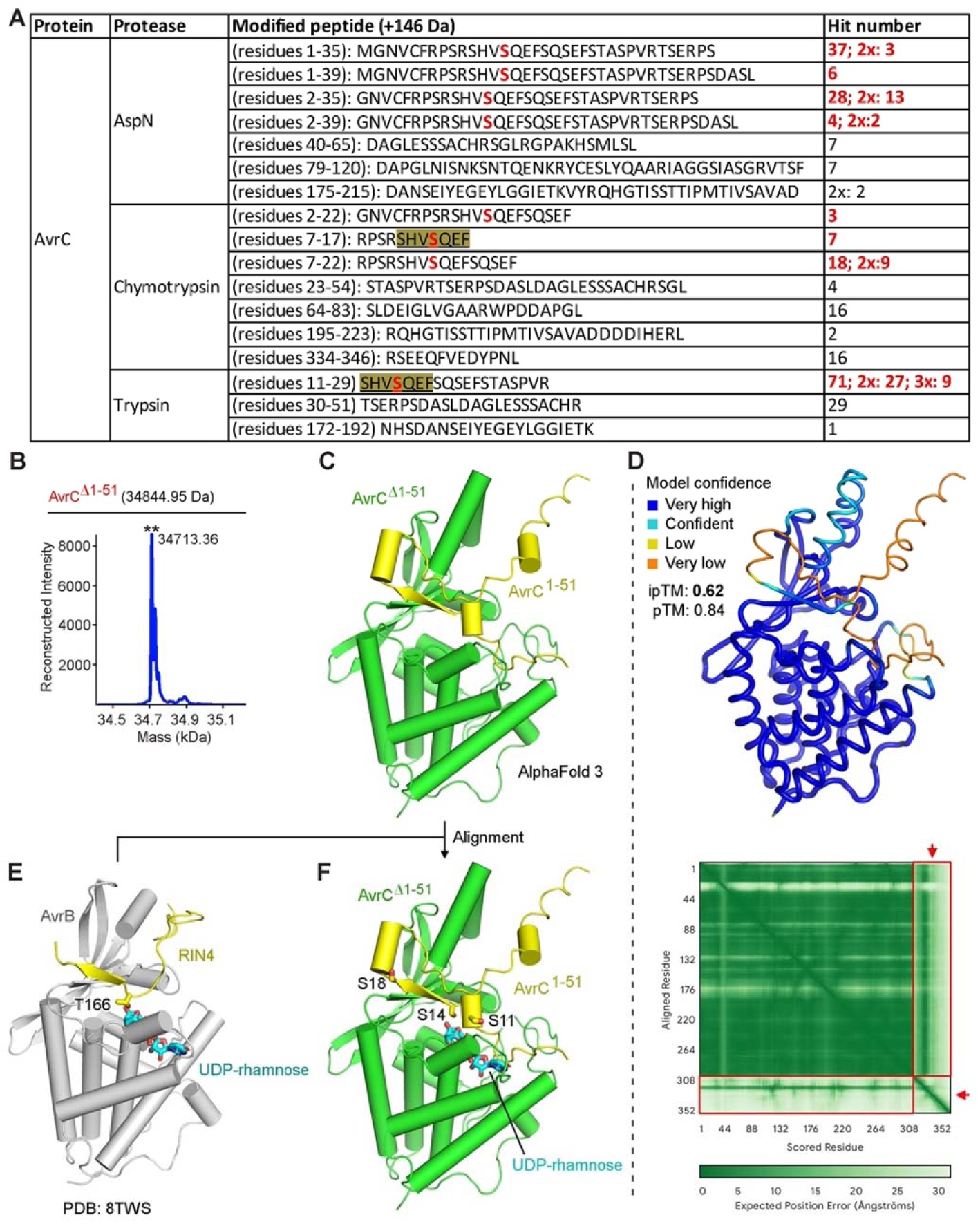
Auto-rhamnosylation of AvrC. (A) PTM analysis of AvrC using proteases AspN, chymotrypsin, or trypsin. Peptides identified with modification of +146 Da are listed. Minimum overlapping region with modification is highlighted. S14 is indicated in red. Peptides carrying 2 or 3 modification sites are indicated with 2x or 3x. (B) Intact mass analysis of AvrC^Δ1-51^ purified from BL21 (DE3). “**” indicates the peak likely caused by loss of initiator methionine (−131 Da). (C) Predicted AlphaFold structure model of the complex of AvrC^Δ1-51^ and AvrC^1-51^. (D) Model confidence for AvrC^Δ1-51^-AvrC^1-51^ complex in (C). Red arrows and boxes indicate interaction prediction between AvrC^Δ1-51^ and AvrC^1-51^. (E) Structure of AvrB bound with RIN4 peptide and UDP-rhamnose (PDB code: 8TWS). (F) Superimposition of the predicted AvrC^Δ1-51^-AvrC^1-51^ complex in (C) and the AvrB-RIN4-UDP-rhamnose complex in (E).

We next assessed the catalytic activity of AvrC *in vitro* using a rhamnosylation assay. For this evaluation we wanted to determine which diphosphate nucleotide AvrC favors. In the thermal shift assay (or differential scanning fluorimetry) using a 384-well system, we observed AvrC, like AvrB, interacted with multiple nucleotides, with UDP showing the most significant binding (Fig. S2, S3) (11). We therefore used UDP-rhamnose as the potential co-substrate for AvrC in our rhamnosylation assay. Using GST-AvrC^1-51^ as the protein substrate and the truncated AvrC^Δ1-51^ as the enzyme, protein intact mass analysis showed that GST-AvrC^1-51^ was rhamnosylated by AvrC^Δ1-51^ (Fig. 2A).

**Fig. 2.**
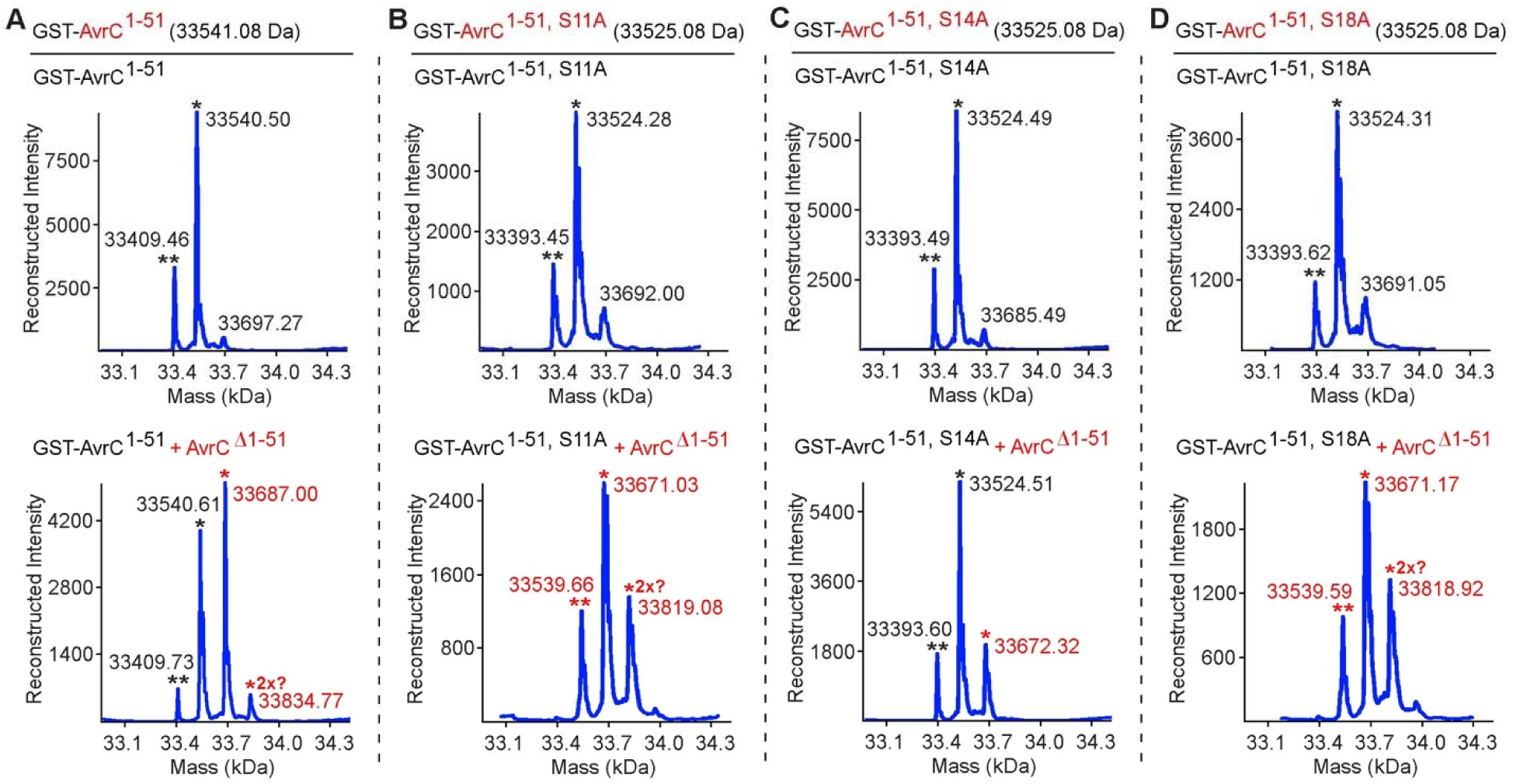
Rhamnosylation of AvrC^1-51^ by AvrC^Δ1-51^. (A-D) Intact mass analysis of GST-AvrC^1-51^ (WT, S11A, S14A, and S18A) after incubation with AvrC^Δ1-51^ in a rhamnosylation assay. “*” symbols in black indicate mass peaks close to the expected mass and “**” symbols in black indicate peaks likely caused by loss of initiator methionine (−131 Da). “*” or “**” symbols in red indicate rhamnosylation peaks with a shift of +146 Da (or likely 2× +146 Da).

To determine if the rhamnosylation site was S14 as predicted by AlphaFold, we made AvrC^Δ1-51^ mutants of serine residues (S11A, S14A, and S18A) near or at the predicted rhamnosylated site. S14A mutation, but not S11A or S18A, greatly repressed the rhamnosylation of AvrC^Δ1-51^ (Fig. 2B-D). Interestingly, the isolated GST-AvrC^1-51^ constructs could likely be rhamnosylated at one additional site by AvrC^Δ1-51^ (Fig. 2). However, no additional rhamnosylation product of the full-length mutant AvrC^S14A^ was observed in the intact mass analysis (Fig. S4). Therefore, we conclude that S14 is the dominant auto-rhamnosylation site in the full-length AvrC protein.

### The core Fido domain of AvrC can rhamnosylate plant protein RIN4

AvrB rhamnosylates the plant “guardee” protein RIN4 (11). However, no substrate(s) for AvrC from host cells has been identified thus far. To identify potential AvrC substrate(s), we performed a pull-down assay for AvrC using *A. thaliana* leaf lysate, followed by mass spectrometry. Proteins enriched in mass spectrometry were also screened for potential interaction with AvrC using AlphaFold complex prediction (Fig. S5A). As expected, this method successfully identified *A. thaliana* RIN4 as the binding protein of AvrB with high interaction scores (ipTM≥0.8) from the AlphaFold complex prediction (Fig. S5B). While none of the AvrC hits were predicted to bind to AvrC with an interaction score above the threshold (ipTM≥0.6), enrichment scores of RIN4 for AvrC in mass spectrometry were in the same magnitude as those for AvrB, likely indicating an interaction between RIN4 and AvrC (Fig. S5B). We therefore predicted the interaction using shorter RIN4 constructs (Fig. S6). The ipTM scores for these AvrC-RIN4 complexes were not as high as those for the AvrC^1-51^ with AvrC^Δ1-51^ complex or the AvrC^11-18^ with AvrC^Δ1-51^ complex, but we noticed that the short β-strands containing T166 and T21 in RIN4 show potential for interaction with AvrC (Fig. S6).

We then tested if *A. thaliana* RIN4 could interact with and be rhamnosylated by AvrC. In the pull-down assay, we observed an interaction of RIN4 with AvrC and AvrC^Δ1-51^, despite observing no interaction between AvrC^1-51^ and AvrC^Δ1-51^ (Fig. S7A). Surprisingly, RIN4 protein co-expressed with full-length AvrC in BL21 (DE3) was not rhamnosylated (Fig. S7B, S8A). We wondered if the unstructured and flexible N-terminus of AvrC prevented RIN4 rhamnosylation. To test this hypothesis, we used AvrC core Fido domain without the N-terminus (AvrC^Δ1-51^) in the assay with RIN4 for rhamnosylation (Fig. 3). Indeed, AvrC^Δ1-51^ modified GST-RIN4 in the rhamnosylation assay to the same extent as what had been observed for AvrB (Fig. 3A) (11). AlphaFold predictions suggest T21 and T166 residues of RIN4 may be modified by AvrC (Fig. S6). In our *in vitro* assay using AvrC^Δ1-51^, we only observed rhamnosylation of GST-RIN4^149-211^, but not GST-RIN4^1-148^ (Fig. 3B, 3C), suggesting T166, but not T21 may be modified. Indeed, we observed the mutant GST-RIN4^T166A^ was not rhamnosylated by AvrC^Δ1-51^ (Fig. 3D). Rhamnosylation of RIN4 by either AvrB or AvrC^Δ1-51^ was detected by our RIN4^T166-Rha^ antibody (Fig. 3E). Hence, T166 of *A. thaliana* RIN4 is the rhamnosylation site for both AvrB and AvrC.

**Fig. 3.**
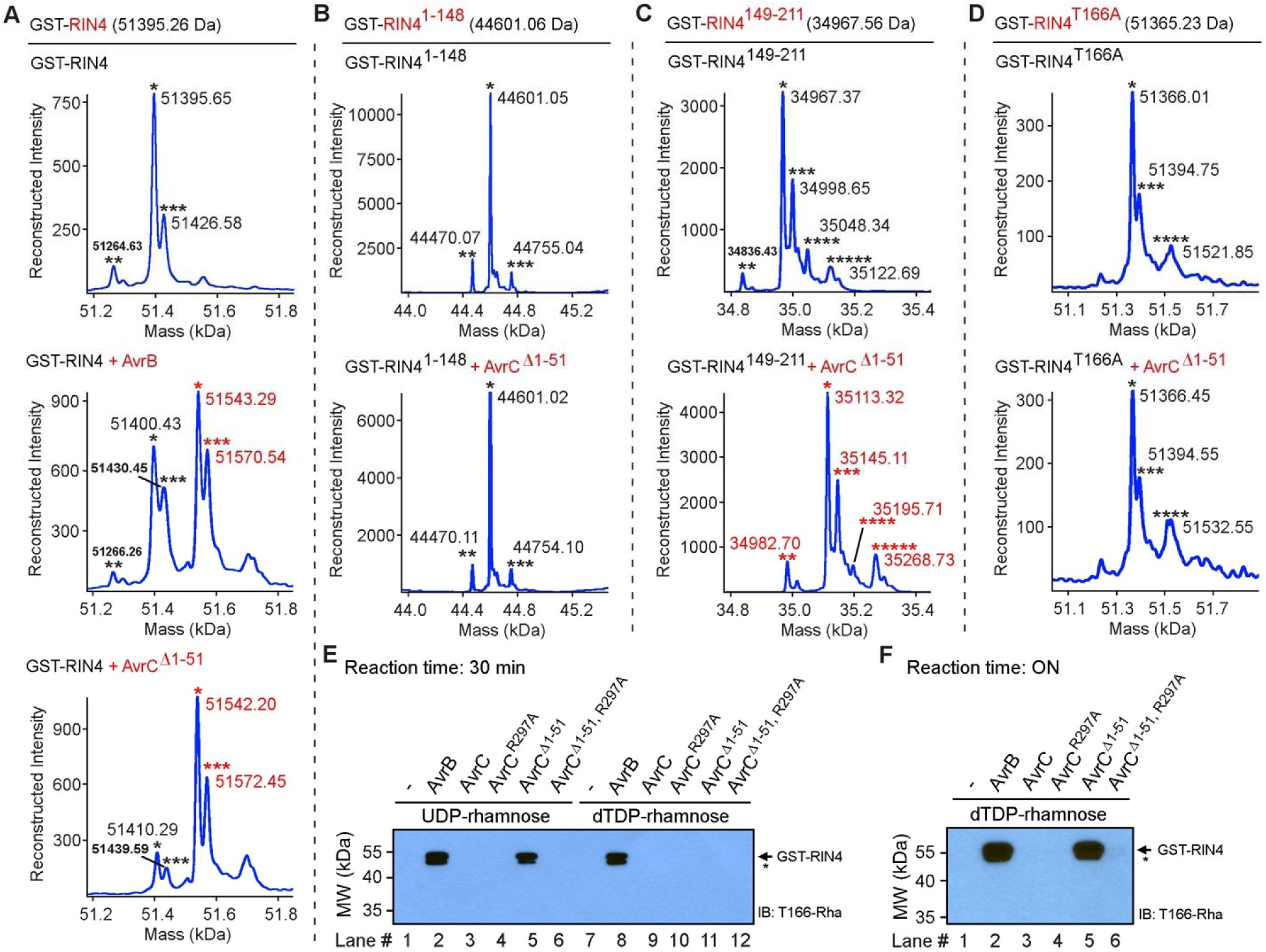
Rhamnosylation of RIN4 by AvrC^Δ1-51^. (A) Intact mass analysis of GST-RIN4 after incubation with AvrB or AvrC^Δ1-51^ in a rhamnosylation assay. “*” symbols in black indicate mass peaks close to the expected mass. “**” symbols in black indicate peaks likely caused by loss of initiator methionine (−131 Da). “***” symbols in black indicate peaks with unknown modification (∼ +30 Da). “*” or “***” symbols in red indicate rhamnosylation peaks with a shift of +146 Da. (B) Intact mass analysis of GST-RIN4^1-148^ after incubation with AvrC^Δ1-51^ in a rhamnosylation assay. “*” symbols in black indicate mass peaks close to the expected mass. “**” symbols in black indicate peaks likely caused by loss of initiator methionine (−131 Da). “***” symbols in black indicate peaks with unknown modification (∼ +54 Da). (C) Intact mass analysis of GST-RIN4^149-211^ after incubation with AvrC^Δ1-51^ in a rhamnosylation assay. “*” symbols in black indicate mass peaks close to the expected mass. “**” symbols in black indicate peaks likely caused by loss of initiator methionine (−131 Da). “***”, “****”, or “*****” symbols in black indicate peaks with unknown modifications. Red “*”, “**”, “***”, “****”, or “*****”symbols indicate rhamnosylation peaks with a shift of +146 Da. (D) Intact mass analysis of GST-RIN4^T166A^ after incubation with AvrC^Δ1-51^ in a rhamnosylation assay. “*” symbols in black indicate mass peaks close to the expected mass. “***” and “****” symbols in black indicate peaks with unknown modifications (∼ +54 Da). (E, F) Rhamnosylation of RIN4 by AvrB or AvrC with UDP-rhamnose or dTDP-rhamnose as a co-substrate. Rhamnosylated RIN4 was detected by the RIN4^T166-Rha^ antibody. The reactions were performed for 30 min or overnight (ON). “*” indicates likely degraded protein.

Auto-rhamnosylation of the AvrC protein purified from BL21 (DE3) likely results from bacterial dTDP-rhamnose serving as the intracellular co-substrate instead of UDP-rhamnose commonly found in plants (11, 16). We then tested if dTDP-rhamnose can be used as a co-substrate for AvrC in the rhamnosylation of RIN4. As expected, we found dTDP-rhamnose to be a co-substrate for AvrC upon overnight incubation (Fig. 3F), albeit less favorable than UDP-rhamnose, which required a shorter 30-minute incubation (Fig. 3E, 3F). The presence of AvrC N-terminus inhibited RIN4 rhamnosylation (Fig. 3E, 3F, S8B). This inhibition was not abolished by S14A or S14F mutant of AvrC (Fig. S8B). A truncation construct, AvrC^Δ6-30^, that lacks the region predicted to interact with AvrC^Δ1-51^ (Fig. 1D) also failed to rhamnosylate RIN4 (Fig. S8B). Therefore, the N-terminus of AvrC likely inhibits rhamnosylation of RIN4 through a yet to be identified mechanism. Surprisingly, RIN4 co-expressed with AvrC^Δ1-51^ in BL21 (DE3) was not rhamnosylated (Fig. S8A).

### AvrC shows broad substrate specificity

Despite showing a poor AlphaFold interaction score with AvrC (ipTM=0.17), we noticed that *A. thaliana* coatomer (or coat protein complex I, COPI) subunit epsilon-1 (COPE1 or ε-1-COP) exhibited mass spectrometry enrichment scores comparable to RIN4 in the AvrC pull-down assay (Fig. S5). We tested if *A. thaliana* COPE1 could be an alternative substrate for AvrC. Using a 6xHis-COPE1 construct, we purified COPE1 protein and obtained two species, one being full-length (1–292) and the other being truncated (1–274) (Fig. 4A, S9). Like RIN4, COPE1 full-length and truncated proteins co-expressed with AvrC were not rhamnosylated (Fig. S9). In the *in vitro* rhamnosylation assay, AvrB did not modify COPE1^1-292^ or COPE1^1-274^ (Fig. 4A). However, COPE1^1-292^ and COPE1^1-274^ could both be modified by AvrC^Δ1-51^ (Fig. 4A). COPE1 has a C-terminal *α*-helix that includes residues 275-292 (Fig. 4B). COPE1^1-274^ being a better *in vitro* substrate than the full-length (Fig. 4A) indicates that a residue near the C-terminal *α*-helix might be the rhamnosylation site that is exposed after truncation. Consistent with these findings, PTM analysis of rhamnosylated COPE1^1-292^ and COPE1^1-274^ proteins suggest residues H252-Y260 most likely carry the rhamnosylation site for AvrC^Δ1-51^ (Fig. S10). We then tested all serine/threonine mutants (S256A, S257A, and S258A) within H252-Y260 (Fig. 4C-4E). Only S258A was observed to completely abolish rhamnosylation, suggesting that S258 is likely the rhamnosylation site on COPE1 for AvrC^Δ1-51^.

**Fig. 4.**
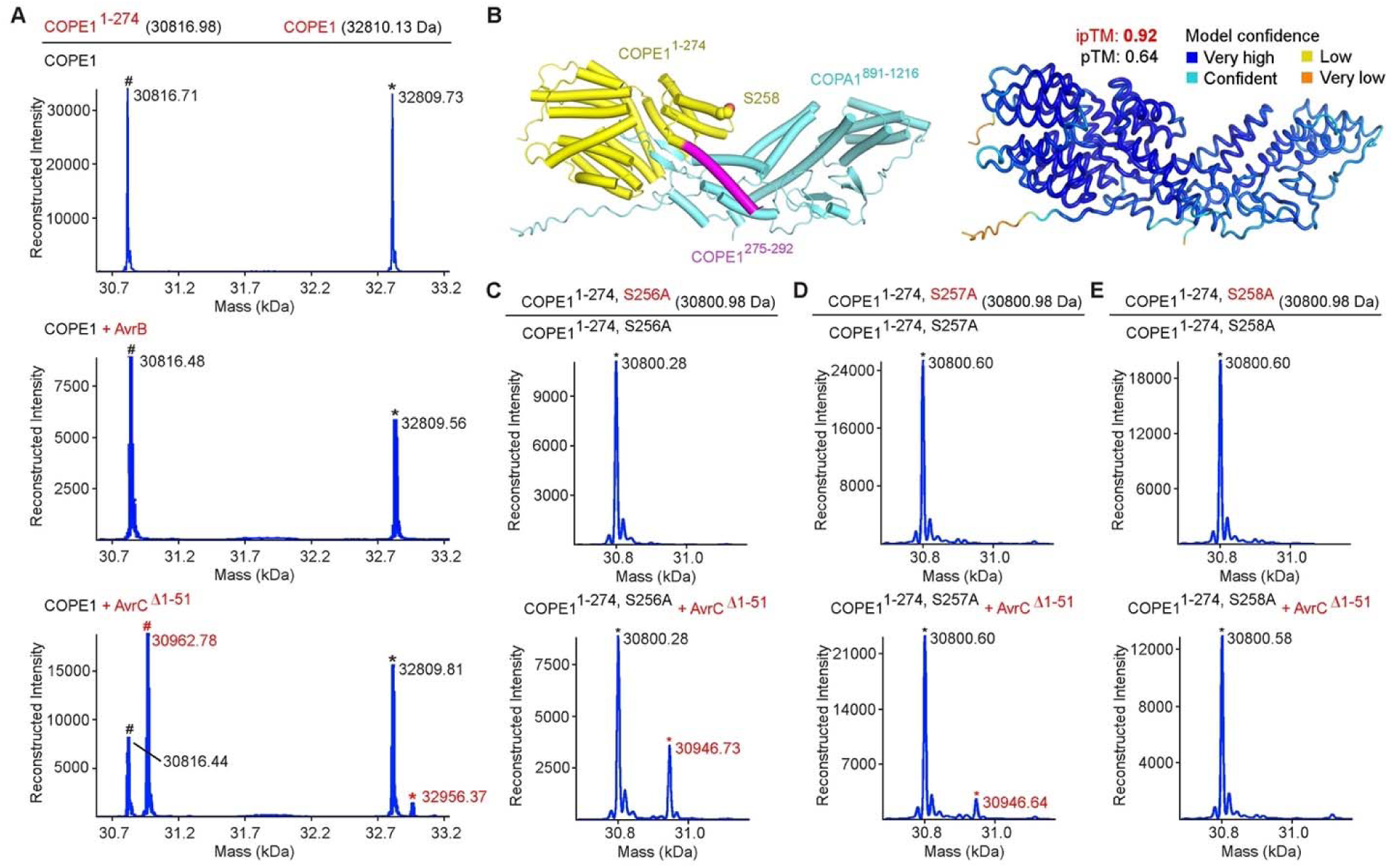
Rhamnosylation of COPE1 by AvrC^Δ1-51^. (A) Intact mass analysis of COPE1 after incubation with AvrC^Δ1-51^ in a rhamnosylation assay. “*” symbols in black indicate mass peaks close to the expected mass. “#” symbols in black indicate potential degradation products which contain residues 1-274. “*” or “#” symbols in red indicate rhamnosylated peaks with a shift of +146 Da. (B) Predicted AlphaFold structure model of COPE1-COPA1 complex. 1-274 of COPE1 is shown in yellow and 275-292 is shown in magenta. S258 is shown as spheres. Model confidence image is shown on the right. (C-E) Intact mass analysis of COPE1^1-274, S256A^, COPE1^1-274, S257A^, and COPE1^1-274, S258A^ after incubation with AvrC^Δ1-51^ in a rhamnosylation assay. “*” symbols in black indicate mass peaks close to the expected mass. “*” symbols in red indicate rhamnosylation peaks with a shift of +146 Da.

To test the interaction between COPE1 and AvrC, we made a GST-COPE1-6xHis construct and purified the protein using affinity resin against either the GST or the 6xHis tag (Fig. S11). Surprisingly, we noticed that the purified protein using the 6xHis tag contained two major species, whereas only one major species was observed using the GST tag for purification. This difference was due to a truncation within the GST tag and not COPE1, supported by the observation that the truncated fragment was only detected by the GST antibody, but not the His antibody after removal of COPE1-6xHis (Fig. S11). We then made a GST-COPE1^1-274^ construct without a C-terminal 6xHis tag, as COPE1^1-274^ was a better *in vitro* substrate for AvrC than the full-length. Interestingly, the pull-down assay indicated that AvrC non-specifically interacted with a non-related human protein (with a GST tag), which was adopted as a negative control (Fig. S12). However, neither GST-COPE1-6xHis nor GST-COPE1^1-274^ interacted with AvrC or AvrC^Δ1-51^, suggesting a poor interaction between AvrC and COPE1 (Fig. S12).

As expected, rhamnosylation of COPE1-6xHis by AvrC was observed by intact mass analysis using the cleaved GST-COPE1-6xHis sample (Fig. S13A). Intact mass analysis of the truncated GST tag protein in the sample suggested that an alternative start codon in GST at L100 was used, resulting in a protein of GST^Δ1-99, L100M^ (Fig. S13B). Interestingly, the truncated GST was rhamnosylated by AvrC^Δ1-51^ while the complete GST tag was not (Fig. S13B), likely due to the exposure of a rhamnosylation site after truncation.

The observations above suggest AvrC non-specifically binds to and modifies multiple proteins. We continued to assess the broad substrate specificity of AvrC by analyzing the *A. thaliana* COPI complex that is responsible for vesicle transport from Golgi to endoplasmic reticulum (17, 18). Using AlphaFold to predict if AvrC interacted with other *A. thaliana* COPI subunits, we observed two subunits showed good interaction scores: COPZ1 (or ζ-1-COP) (ipTM_1_=0.77, ipTM_2_=0.77) and COPD (or -COP) (ipTM_1_=0.62, ipTM_2_=0.5). As *A. thaliana* COPD was not expressed well, we only tested *A. thaliana* COPZ1 using pull-down and rhamnosylation assays. In the pull-down assay, *A. thaliana* COPZ1 showed an interaction with AvrC and AvrC^Δ1-51^ (Fig. S14). Human COPI subunits were also tested in our AlphaFold complex predictions and only COPZ2 showed good confidence scores (ipTM_1_=0.64, ipTM_2_=0.72). Human COPZ2 interacted with both AvrC^Δ1-51^ and AvrC as well in the pull-down assays (Fig. S14). *In vitro* rhamnosylation assay indicated *A. thaliana* COPZ1, but not human COPZ2, was modified by AvrC^Δ1-51^ (Fig. S15A, S15B). Our PTM mass spectrometry analysis confirmed the rhamnosylation of *A. thaliana* COPZ1 by AvrC^Δ1-51^ (Fig. S15C).

### Activity of AvrB and AvrC in *N. benthamiana* leaves

To compare the hypersensitive response symptoms caused by AvrB and AvrC, we tested the two effectors using the *N. benthamiana* model system. *N. benthamiana* encodes a RIN4 homologue (Nb-RIN4) with 77% similarity to *A. thaliana* RIN4 (At-RIN4) over residues 147-178 of At-RIN4 that mediate binding to effectors, including the rhamnosylation site T166 (Fig. S16A). As expected, recombinant Nb-RIN4 was rhamnosylated by AvrB and AvrC^Δ1-51^ using *in vitro* rhamnosylation assays (Fig. S16B) (11). Using infiltration assays on *N. benthamiana* leaves, AvrB was able to induce symptoms of hypersensitive response even though the endogenous Nb-RIN4 protein could not be detected (Fig. S17, S18). At-RIN4 or Nb-RIN4 co-expressed with AvrB in *N. benthamiana* leaves was not detected in the absence or presence of a P19 RNA silencing suppressor that increases protein expression level (Fig. S17, S18). Interestingly, higher expression level of AvrB likely induced rhamnosylation of a high molecular weight, unknown factor that was recognized by the RIN4^T166-Rha^ antibody (Fig. S17B, S18B). This additional factor may be related to the symptoms caused by AvrB.

Unlike AvrB, neither AvrC nor AvrC^Δ1-51^ caused obvious symptoms in the *N. benthamiana* model system and displayed undetectable protein expression (Fig. S19A, S19B). Glycine myristoylation-mediated association with membranes likely regulates the effector function of AvrB and AvrC (19, 20). Therefore, we tested a few AvrB and AvrC constructs with and without the myristoylation site (G2 in Fig. S1A). Recombinant AvrB^Δ1-26^ protein lacking the myristoylation site and AvrB^Δ10-26^ protein with this site were functional in the *in vitro* rhamnosylation assay (Fig. S19C). However, a swap of the AvrB N-terminus onto AvrC core Fido domain (AvrB^1-9^-AvrC^Δ1-51^) was catalytically inactive (Fig. S19C), similar to the inhibition by AvrC N-terminus mentioned above. In multiple infiltration assays, AvrB^Δ1-26^ and AvrB^Δ10-26^ could induce disease symptoms (Fig. S19A, S19B). However, AvrC^Δ6-30^ showed no symptoms in these assays and exhibited undetectable protein expression. The AvrB^1-9^-AvrC^Δ1-51^ construct showed no or weak symptoms although the protein expression was observed (Fig. S19A, S19B).

## Discussion

We previously showed that the prototypical *P. syringae* avirulence effector AvrB is a rhamnosyltransferase in the AvrB-RIN4-RPM1 axis of plant-pathogen interaction (11). Here we show that the related fido domain-containing effector AvrC also functions as a rhamnosyltransferase. While AvrB showed no auto-rhamnosylation activity (11), auto-rhamnosylation at S14 was observed on the recombinant protein of full-length AvrC. The core Fido domain (AvrC^Δ1-51^) lacking the N-terminus could bind to and rhamnosylate the plant “guardee” protein RIN4. Host vesicle transport pathways have also been shown to be targeted by bacterial effectors (21–23). We found that AvrC^Δ1-51^ could *in vitro* rhamnosylate *A. thaliana* vesicle transport coatomer subunits (COPE1 and COPZ1) and even a truncated GST protein GST^Δ1-99, L100M^.

As a promiscuous rhamnosyltransferase, AvrC likely binds to and modifies various factors when delivered into host cells. These potential non-specific interactions and modifications may be involved in the biological functions of AvrC. Furthermore, AvrC may have additional host specific targets found in soybean cultivars, as AvrC induces disease phenotypes in this host (5, 15). Future studies are required to address these questions about AvrC virulence and/or avirulence activities.

In contrast to truncated AvrC^Δ1-51^, full-length AvrC (WT, S14A, or S14F) did not modify RIN4 *in vitro*. AvrC^Δ6-30^ and AvrB^1-9^-AvrC^Δ1-51^, which retain a host myristylation targeting site, could not rhamnosylate RIN4, either. These results indicate that residues proceeding AvrC core Fido domain likely regulate the enzymatic activity through an unknown mechanism. Bacterial effectors can be activated by a host component (e.g., IP6) or modification by another effector (e.g., *Legionella pneumophila* metaeffectors) (24–26). When delivered into host cells, AvrC will likely be modified by host enzyme(s), e.g., myristoylation as has been observed for related bacterial effector proteins (19, 20). The N-terminus of AvrC may also be cleaved. These events could contribute to activating the rhamnosyltransferase and causing virulence/avirulence phenotypes in its target soybean cultivars. AvrB and AvrC are independent effectors (5, 14, 15), but AvrC occurs on the same plasmid with effector AvrD in a few *P. syringae* strains and AvrC can suppress the avirulence activity of another effector AvrPphF (27, 28). Therefore, whether and how AvrC interacts host factors and/or other bacterial effectors remains to be studied.

In summary, our work shows that combination of biochemistry, mass spectrometry, and AlphaFold prediction tools can contribute to identification of interacting proteins and modification sites for AvrC. The findings provide valuable insights into future studies on other Fido enzymes. Our observations indicate that AvrC is a rhamnosyltransferase with broad substrate specificity. The reduced specificity of AvrC may be utilized to produce rhamnosylated proteins or peptides that may have potential applications, e.g., synthesizing rhamnose-conjugated antibodies for complement-dependent targeted cancer cell killing through recruiting natural anti-rhamnose antibodies (29). AvrC or AvrB may also be used as a scaffold for protein engineering/design to construct glycosyltransferases with desired specificities in the age of artificial intelligence advancement (30).

## Experimental procedures

### Protein expression and purification

The cDNA encoding *P. syringae* AvrC was cloned into the pET-29b vector with a C-terminal 6xHis tag. The cDNA encoding *A. thaliana* RIN4, COPE1, COPZ1, or human COPZ2 was cloned into a modified pET-15b vector (with an N-terminal 6xHis tag followed by a DrICE protease cutting site) or a modified pGEX-4T-2 vector with/without a C-terminal 6xHis tag.

The plasmid was transformed into *E. coli* strain BL21 (DE3) for protein expression. For co-expression, AvrC and RIN4/COPE1 plasmids were co-transformed into BL21 (DE3). The bacteria were cultured at 37. When OD_600_ reached ∼1.0, the temperature was adjusted to 22, and 0.2 mM IPTG was added for overnight induction. Cells were collected by centrifugation, re-suspended in lysis buffer B1 (25 mM Tris-HCl (pH 8.0) and 150 mM NaCl). After cell disruption and removal of cell debris by centrifugation at 22,000 g for 1 h, the supernatant was loaded to Ni^2+^-NTA resin (Qiagen). The resin was washed by buffer B2 (lysis buffer B1 with 300 mM NaCl), and sequentially by buffer B3 (lysis buffer B1 containing 10 mM imidazole). Protein bound to the resin was eluted by buffer B4 (lysis buffer B1 containing 250 mM imidazole). The eluted protein was dialyzed against buffer B5 (25 mM HEPES (pH 7.4) and 150 mM NaCl) with 7 kDa or 10 kDa Slide-A-Lyzer Dialysis Cassettes (Thermo Fisher). The protein was flash-frozen by liquid nitrogen and stored at −80 for later use.

### Pull-down assay

10 µg of bait protein (with GST tag) and 10 µg of prey protein were mixed with 10 µL of Glutathione Agarose resin in 500 µL of pull-down buffer containing 5 mg of BSA for blocking. Pull-down buffer was composed of 25 mM HEPES (pH7.4), 150 mM NaCl, 1 mM DTT, and 0.1% Triton X-100. Samples were incubated at 4 for 30 min. After centrifugation at 1,000 g for 1 min, the supernatant was discarded. The resin was then washed 3 times with 500 µL of pull-down buffer. SDS sample buffer was mixed with the remaining resin for SDS-PAGE and Coomassie Brilliant Blue staining.

### Mass spectrometry

Protein intact mass analysis was performed following the previous protocol (11). Protein LC-MS/MS analysis to determine the modification site was performed similarly as reported previously (11). Because loss of the modification was observed upon fragmentation via higher-energy collision dissociation, various proteases including AspN, chymotrypsin, and trypsin were used to generate multiple peptides for comparison.

### Thermal shift assay

The thermal shift assay was performed similarly as previously reported with modifications (11). Triplicate 18 μL reaction systems contained 0.2 mg/mL of AvrB or AvrC protein (∼5 µM). Reaction buffer contained: 25 mM HEPES (pH 7.4), 100 mM NaCl, 1 mM DTT, 1 mM or 4 mM small molecule, and 1:500 diluted SYPRO Orange Protein Gel Stain (Sigma). Reactions were performed in a 384-well PCR plate with a CFX Opus 384 Real-Time PCR System. Samples were subjected to a gradient of temperature 10∼90 (hold 5 s, increase 0.5, rate 0.5 /s). Fluorescent signals were recorded using FRET channel. Melting curves were normalized for each replicate and plotted in GraphPad Prism software. The temperature at the derivative peak was set as the melting temperature, calculated by Bio-Rad CFX Maestro Software.

### In vitro rhamnosylation assay

Rhamnosylation reactions were carried out in a buffer solution containing 25 mM HEPES (pH 7.4), 150 mM NaCl, and 5 mM DTT. Standard reactions included 0.4 µg of AvrB or AvrC and 0.8 µg of GST-RIN4 in a final volume of 25 µL with 1 mM or 100 µM of UDP-rhamnose (MedChemExpress). Reactions were performed at room temperature for 30 min (unless specifically noted) and stopped by the addition of 25 µL of 2x SDS sample buffer.

### Tobacco leaf infiltration assay

The assay was performed following a protocol with modifications (31). The corresponding cDNA was cloned into a modified pCAMBIA1300 vector. The plasmid was electroporated into *Agrobacterium tumefaciens* (GV3101) cells. *A. tumefaciens* cells carrying the gene of interest were grown overnight in LB medium at 30. Cells were harvested (4000 g, 7 min) and resuspended in the infiltration buffer containing 10 mM MES (pH 6.5), 10 mM MgSO_4_, and 150 μM acetosyringone. Cells were adjusted to OD_600_ of 0.4 ∼ 0.8 and incubated at room temperature for 2 ∼ 3 hours. The bacterial suspension was infiltrated into leaves of *N. benthamiana* with a needleless syringe. Leaves were harvested 5 days later for imaging and immunoblotting.

### Plant leaf sample preparation

Harvested tobacco leaves were ground with liquid nitrogen. 5 times (μL/mg) of 2x SDS sample buffer was added. Samples were heated (>95, 10 min) before SDS-PAGE and immunoblotting.

### Immunoblotting

Samples were applied to SDS-PAGE and transferred to PVDF or nitrocellulose membrane for immunoblotting with corresponding antibodies.

### Structure prediction and analysis

Predicted AlphaFold structure models were analyzed (32). Complex prediction was done using AlphaFold server at https://alphafoldserver.com/ (33). Prediction with ipTM≥0.4 was repeated for a second round. Structure images were prepared with PyMol (The PyMOL Molecular Graphics System, Version 2.4, Schrödinger, LLC.).

## Supporting information

This article contains supporting information.

## Acknowledgments

We thank members of the Orth laboratory for insightful discussions and editing. We thank the proteomics core facility at UT Southwestern Medical Center for assistance with protein intact mass analysis. KO is a WW Caruth, Jr. Biomedical Scholar with an Earl A Forsythe Chair in Biomedical Science.

## Funding

Welch Foundation grant I-1561 (KO)

Once Upon a Time…Foundation (KO)

National Institutes of Health Grant R35 GM134945 (KO)

Research Opportunity Seed Fund (ROSF) from University of Florida Office of Research (JF)

## Conflict of interest

The authors declare that they have no conflicts of interest with the contents of this article.

## Notes

### Competing Interest Statement

The authors have declared no competing interest.

