## supporting information for "*Pseudomonas* effector AvrC is a rhamnosyltransferase with broad substrate specificity"

Nalleli Payne *et al.*

**This supporting information PDF file includes:**

Figure S1 to S19

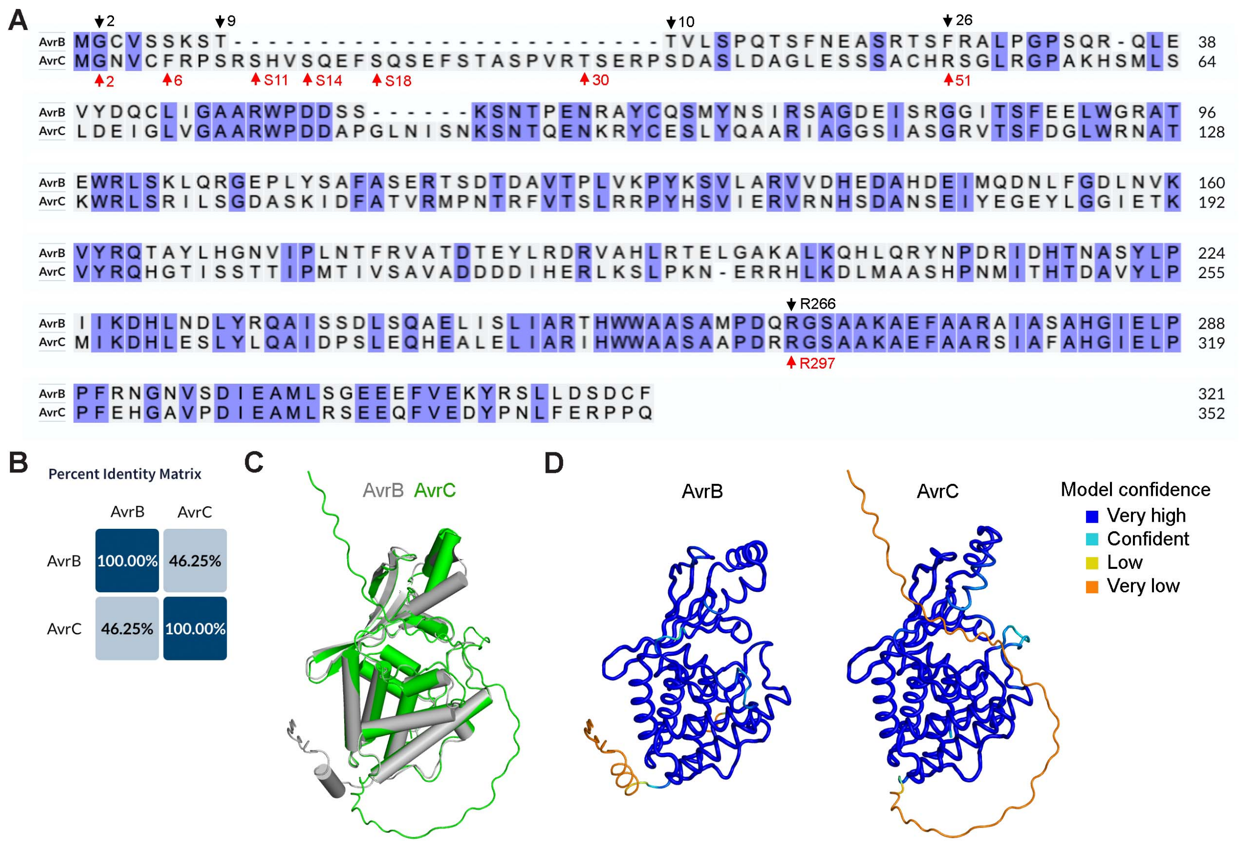


**Figure S1. AvrC is Fido protein similar to AvrB.**

(A) Sequence alignment of AvrB and AvrC done at: https://www.uniprot.org/align. Boundaries and residues of AvrB (black) and AvrC (red) investigated in this study are indicated.

(B) Percent Identity Matrix between AvrB and AvrC obtained at: https://www.uniprot.org/align.

(C) Superimposition of AlphaFold structure models of AvrB (P13835) and AvrC (P13836).

(D) Model confidence for AvrB and AvrC shown in (C).

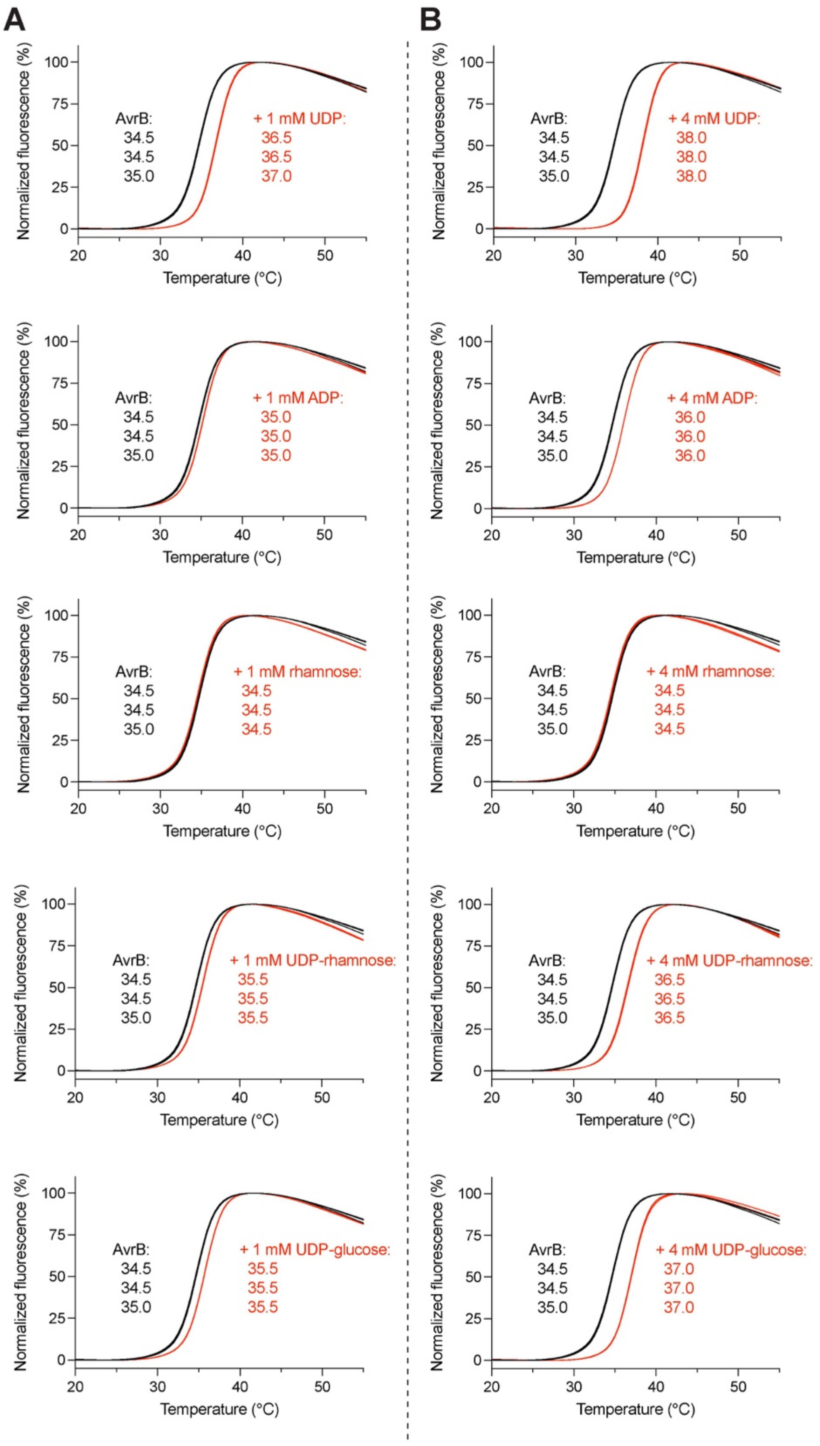


**Figure S2. Thermal shift assay for testing interaction between AvrB and small molecules.**
(A) Curves for AvrB (5 µM) alone and with molecules at a concentration of 1 mM (UDP, ADP, rhamnose, UDP-rhamnose, and UDP-glucose).
(B) Curves obtained with molecules at a concentration of 4 mM.

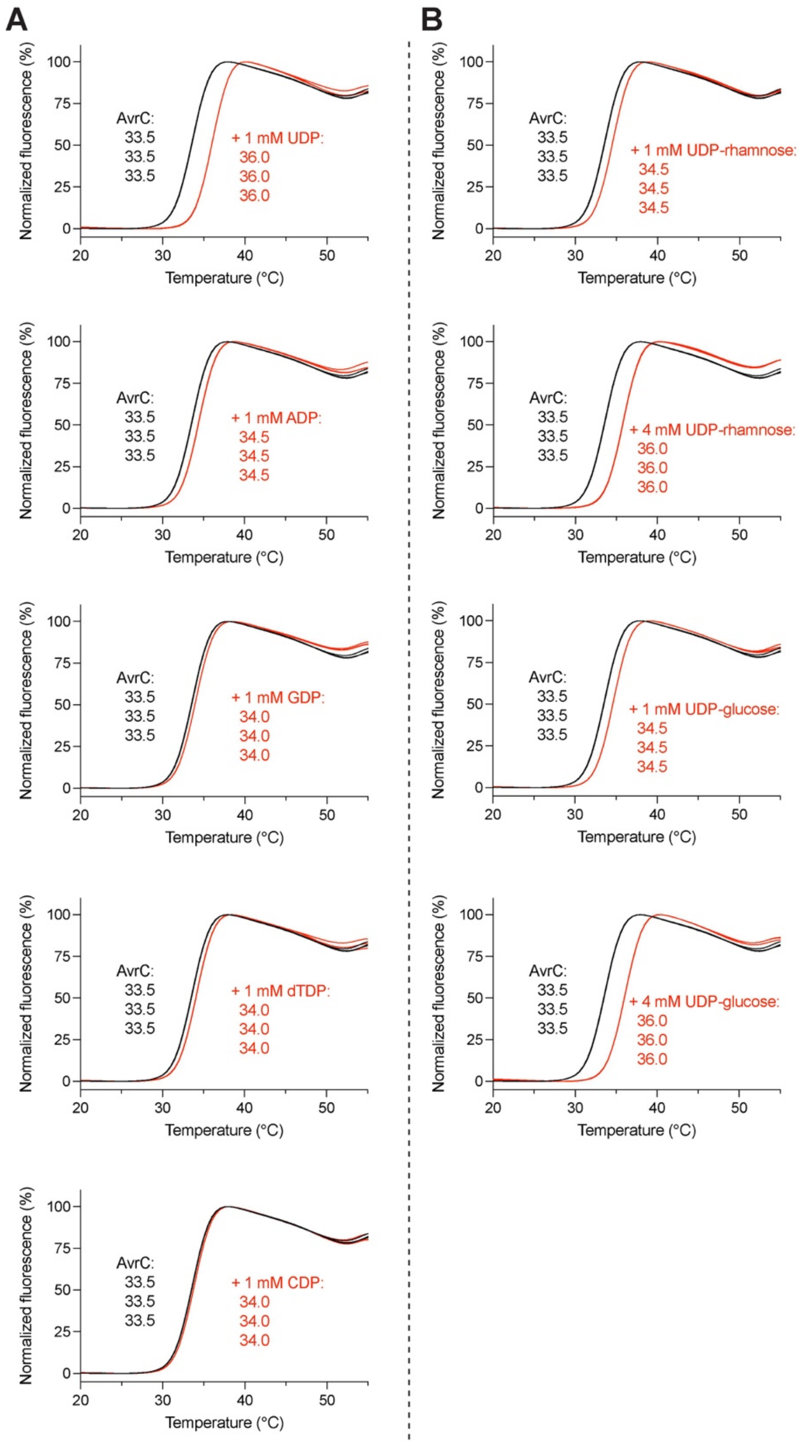


**Figure S3. Thermal shift assay for testing interaction between AvrC and small molecules.**
(A) Curves for AvrC (5 μM) alone and with molecules at 1 mM (UDP, ADP, GDP, dTDP, CDP).
(B) Curves obtained with small molecules at 1 mM and 4 mM (UDP-rhamnose, UDP-glucose).

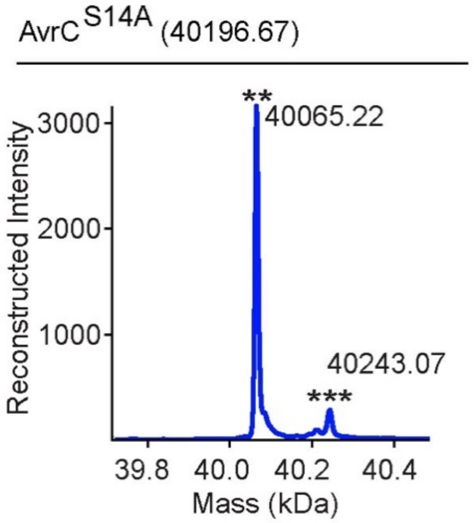


**Figure S4. Intact mass analysis of AvrC$^{S14A}$.**
AvrC$^{S14A}$ purified from BL21 (DE3) was analyzed. "**" indicates the peak likely caused by loss of initiator methionine (-131 Da). "***" indicates the peak likely caused by gluconoylation (+178 Da compared to "**").

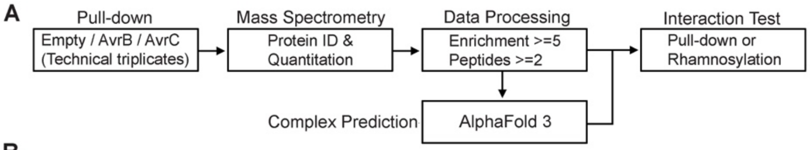


B

| Bait | Prey (Uniprot) | Protein Description | Enrichment 1 | Enrichment 2 | Enrichment 3 | Enrichment Average | ipTM | pTM |
|---|---|---|---|---|---|---|---|---|
| AvrB | Q9T0G1 | Alpha/beta-Hydrolases superfamily protein | 88423488.72 | 44928173.94 | 62109852.43 | 65153838.36 | 0.17 | 0.46 |
| AvrB | Q9CA90 | Glyoxylate/hydroxypyruvate reductase A HPR2 | 11235511.74 | 11220504.78 | 14606767.79 | 12354261.44 | 0.1 | 0.49 |
| AvrB | Q9M1Q0 | RING-type E3 ubiquitin transferase | 26538520.31 | 118.2106306 | 87.21055352 | 8846241.912 | 0.26 | 0.35 |
| AvrB | A0A1I9LMJ4 | **RPM1 interacting protein 4** | **126.6156256** | **150.5419636** | **72.20522898** | **116.4542727** | **0.81**<br>**0.81** | 0.61<br>0.61 |
| AvrC | Q66GP9 | NO-associated protein 1, chloroplastic/mitochondrial | 6.443923981 | 46787546.77 | 61966496.52 | 36251349.91 | 0.34 | 0.49 |
| AvrC | Q8GWK5 | Gibberellin-regulated protein 9 | 25187204.78 | 651.1790265 | 41307328.93 | 22165061.63 | 0.24 | 0.65 |
| AvrC | O80731 | Pectin acetylesterase 3 | 21967755.3 | 24202363.94 | 19106049.01 | 21758722.75 | 0.13 | 0.52 |
| AvrC | Q8VZ31 | Tetratricopeptide repeat (TPR)-like superfamily protein | 8318942.006 | 19867489.48 | 13393343.82 | 13859925.1 | 0.23 | 0.5 |
| AvrC | Q9CA90 | Glyoxylate/hydroxypyruvate reductase A HPR2 | 8360598.037 | 4485109.953 | 12705566.45 | 8517091.479 | 0.1 | 0.48 |
| AvrC | Q9M1Q0 | RING-type E3 ubiquitin transferase | 21934172.79 | 41.52507875 | 70.23192896 | 7311428.182 | 0.26 | 0.36 |
| AvrC | A0A1I9LMJ4 | **RPM1 interacting protein 4** | **347.3787671** | **313.4163964** | **132.3345466** | **264.3765701** | **0.53**<br>**0.49** | 0.56<br>0.57 |
| AvrC | Q9SA78 | **Coatomer subunit epsilon-1** | **147.3433359** | **170.7563348** | **142.6465636** | **153.5820781** | 0.17 | 0.5 |
| AvrC | Q8W0Z9 | Carbon catabolite repressor protein 4 homolog 1 | 17.79451996 | 24.01620073 | 25.56026531 | 22.45699534 | **0.44**<br>**0.4** | 0.59<br>0.6 |
| AvrC | Q1JPL4 | External alternative NAD(P)H-ubiquinone oxidoreductase B1, mitochondrial | 10.88180352 | 18.59927902 | 9.925305103 | 13.13546255 | **0.45**<br>**0.18** | 0.56<br>0.52 |
| AvrC | Q9SY88 | E3 ubiquitin-protein ligase CSU1 | 6.70453251 | 19.28247816 | 8.420445821 | 11.46915216 | **0.44**<br>**0.48** | 0.52<br>0.53 |

**Figure S5. Identification of AvrC-binding proteins from *A. thaliana* leaf lysate.**
(A) Flowchart for identifying proteins that may interact with AvrC.
(B) Proteins that pass through the threshold in mass spectrometry data processing. AlphaFold 3 prediction scores are listed. Candidates with scores of ipTM≥0.4 were predicted for a second round.

**A**

| Protein 1 | Protein 2 (Uniprot / Name) | | ipTM | pTM | Modification Site | |
|---|---|---|---|---|---|---|
| AvrB | A0A1I9LMJ4 | RIN4 | **0.81** | 0.61 | **T166** in Q8GYN5 | |
| | | | **0.81** | 0.61 | **T166** in Q8GYN5 | |
| | Q8GYN5 | RIN4 | **0.81** | 0.62 | **T166** | B |
| | | | **0.81** | 0.63 | **T166** | |
| | | RIN4 (residues 138-177) | **0.81** | 0.86 | **T166** | C |
| | | | **0.82** | 0.86 | **T166** | |
| AvrC$^{\Delta 1\text{-}51}$ | | AvrC (residues 1-51) | **0.62** | 0.84 | **S14** | |
| | | | **0.6** | 0.84 | **S14** | |
| | | AvrC (residues 11-18) | **0.8** | 0.95 | **S14** | D |
| | | | **0.81** | 0.93 | **S14** | |
| AvrC | A0A1I9LMJ4 | RIN4 | 0.53 | 0.56 | **T166** in Q8GYN5 | |
| | | | 0.49 | 0.57 | **T166** in Q8GYN5 | |
| AvrC$^{\Delta 1\text{-}51}$ | Q8GYN5 | RIN4 | 0.55 | 0.62 | **T166** | E |
| | | | 0.56 | 0.62 | **T166** | |
| | | RIN4 (residues 138-177) | 0.34 | 0.86 | | |
| | | | 0.33 | 0.86 | | |
| | | RIN4 (residues 1-130) | 0.55 | 0.71 | **T21** | F |
| | | | 0.56 | 0.71 | **T21** | |
| | | RIN4 (residues 131-211) | 0.41 | 0.77 | T166 | G |
| | | | 0.36 | 0.77 | T166 | |

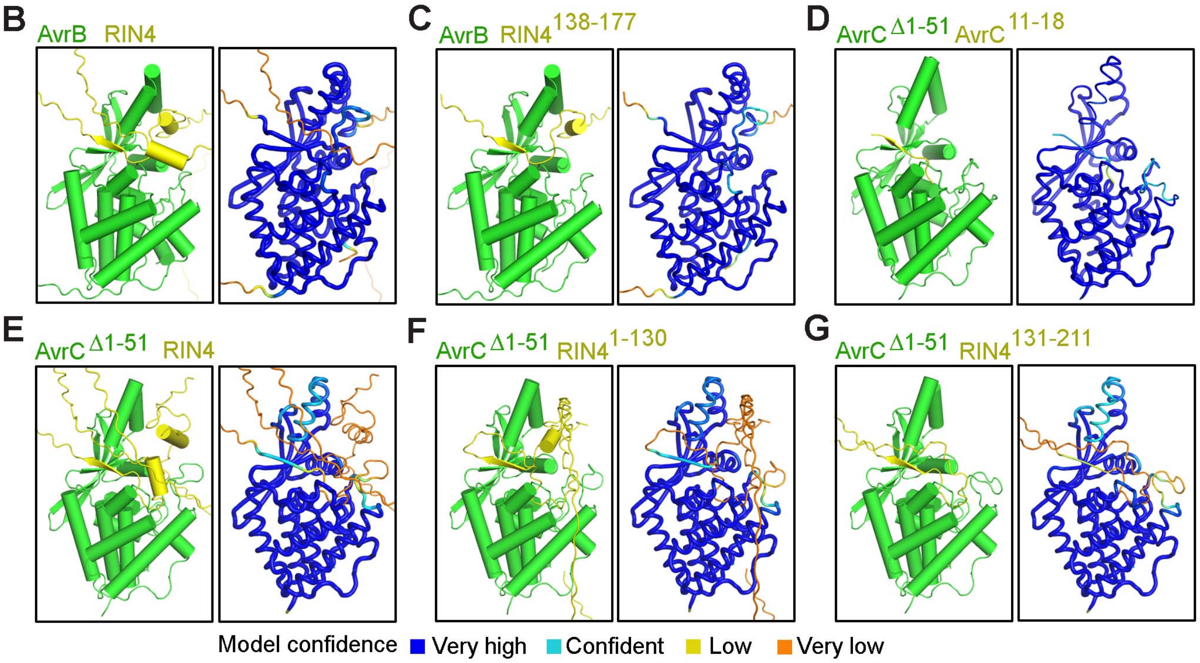


**Figure S6. Predicted rhamnosylation site of RIN4 by AvrC.**
(A) Prediction scores for interaction between AvrB or AvrC and RIN4 or AvrC peptides.
(B-G) Representative prediction models of complexes indicated in (A). Models are shown on the left and prediction confidence images are shown on the right.

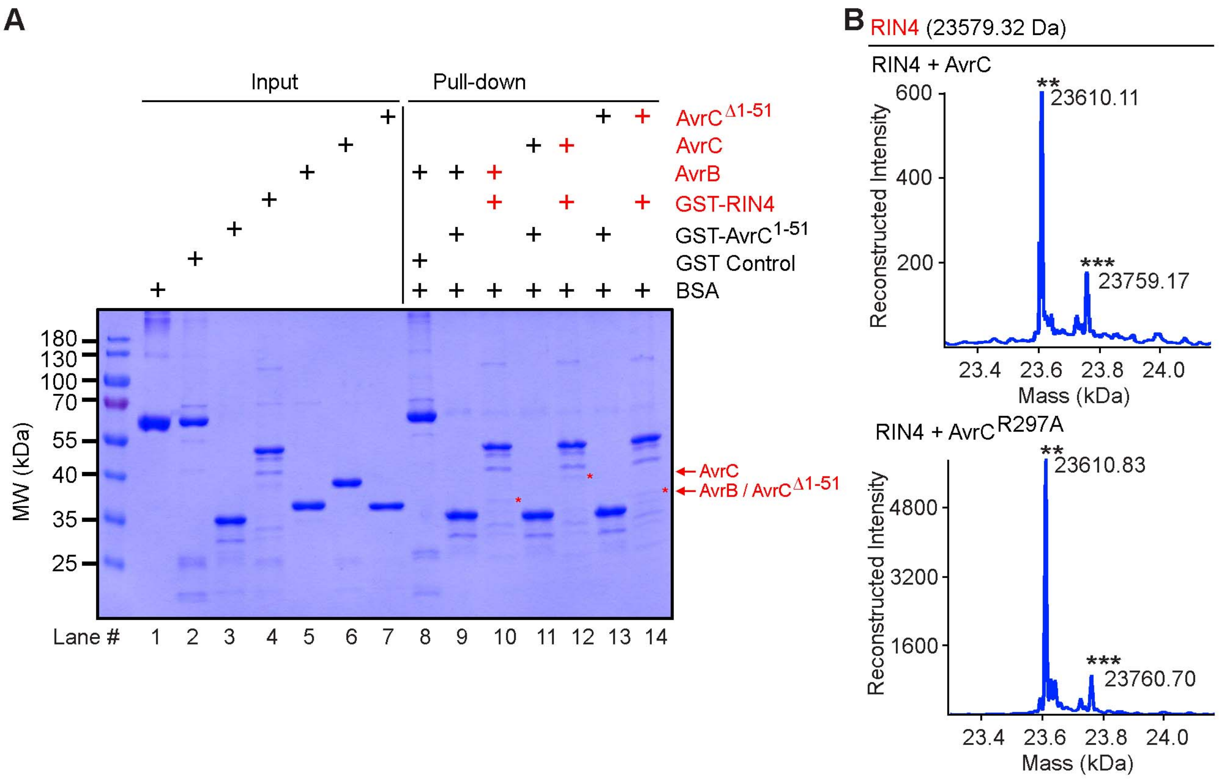


**Figure S7. Interaction between RIN4 and AvrC.**
(A) Pull-down assay for testing interaction between AvrC$^{1-51}$ or RIN4 and AvrC or AvrC$^{\Delta1-51}$. "+" in red indicates a detected interaction. "*" in red indicates a protein that was pulled down.
(B) Intact mass analysis of purified RIN4 proteins that were co-expressed with AvrC (WT and the catalytically dead mutant R297A) in BL21 (DE3). "**" symbols indicate unknown modification peaks (+32 Da). "***" symbols indicate unknown modification peaks (+180 or +181 Da).

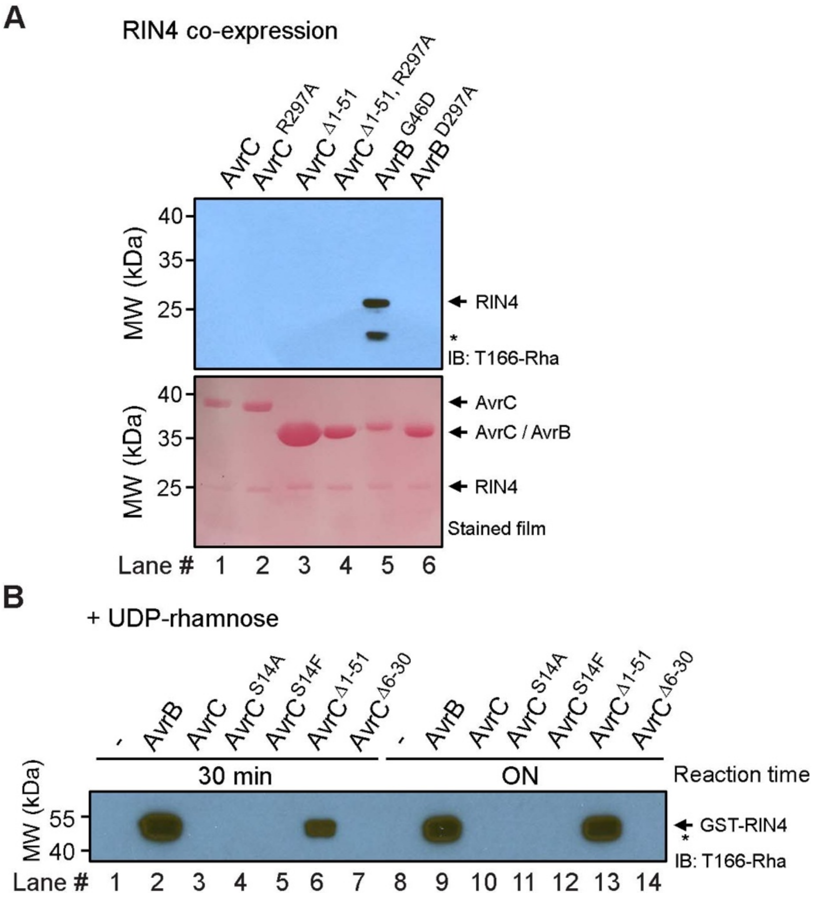


**Figure S8. Rhamnosylation of RIN4 by AvrC.**
(A) Detection of rhamnosylation of purified RIN4 proteins that were co-expressed with AvrC or AvrB in BL21 (DE3). "*" indicates likely degraded protein.
(B) *In vitro* rhamnosylation of RIN4 by AvrC using UDP-rhamnose as the co-substrate. The reactions were performed for 30 min or overnight (ON). "*" indicates likely degraded protein.

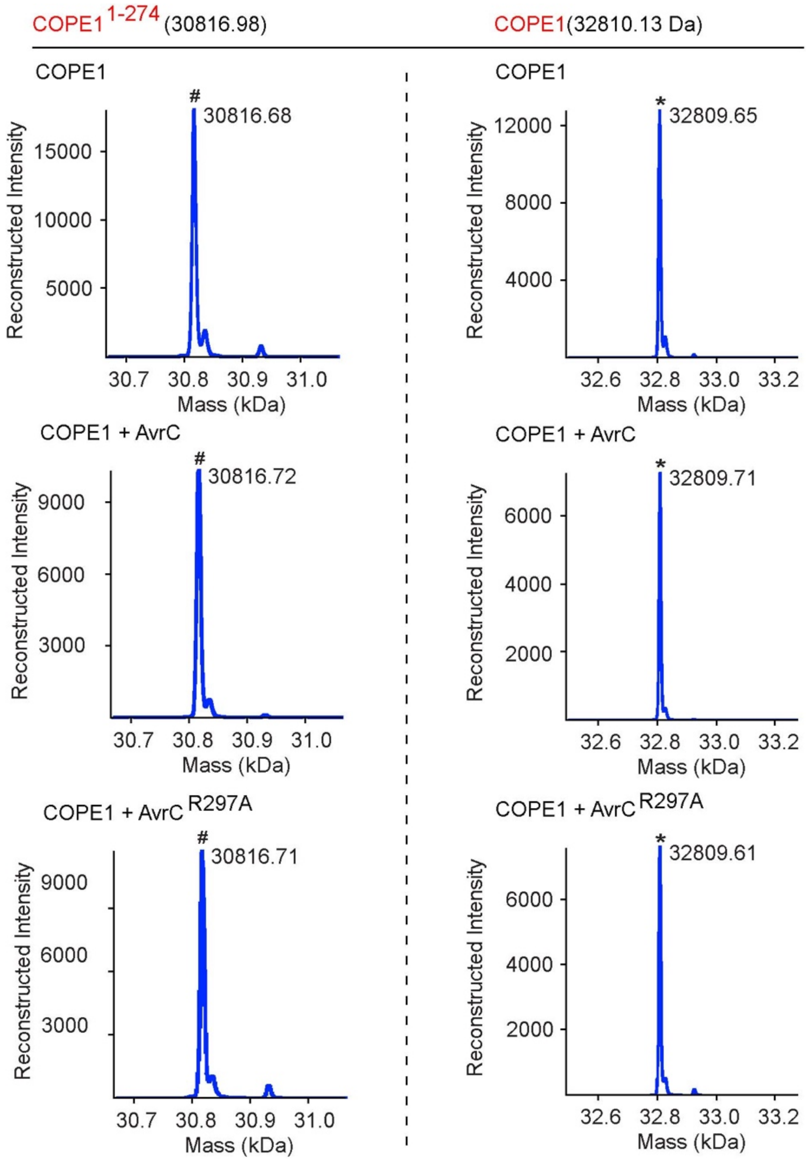


**Figure S9. Intact mass analysis of purified COPE1 proteins.**

Intact mass analysis of purified COPE1 proteins that were expressed alone or with AvrC in BL21 (DE3). "*" symbols indicate peaks close to the expected mass. "#" symbols indicate degradation products which contain residues 1-274.

| Protein | Protease | Modified peptide (+146 Da) | Hit number |
| --- | --- | --- | --- |
| COPE1 | AspN | - | - |
| | Chymotrypsin | (residues 1-10): AH-MASMAGPDHL | 5 |
| | | (residues 99-111): ADPTVGNNAIIRL | 3 |
| | | (residues 179-193): LNLAVGGSKIQEAYL | 1 |
| | | (residues 180-192): NLAVGGSKIQEAY | 1 |
| | | (residues 194-207): IFQDFSEKYPMTSL | 2 |
| | | (residues 252-260): HVGK**SSS**RY | **7** |
| | | (residues 252-261): HVGK**SSS**RYL | **6** |
| | Trypsin | (residues 1-14): AH-MASMAGPDHLFNLR | 2x: 2 |
| | | (residues 88-95): ESTISSLR | 1 |
| | | (residues 96-110): EWLADPTVGNNAIIR | 4; 2x: 3 |
| | | (residues 188-201): IQEAYLIFQDFSEK | 4 |
| | | (residues 277-286): AASAEDNFER | 4 |
| COPE1$^{1-274}$ | AspN | (residues 122-136): DYNEALKHTHSGGTM | 3 |
| | Chymotrypsin | (residues 1-10): AH-MASMAGPDHL | 5 |
| | | (residues 57-71): QLVISEIDEAAATPL | 2 |
| | | (residues 99-111): ADPTVGNNAIIRL | 2 |
| | | (residues 252-260): HVGK**SSS**RY | **91** |
| | | (residues 252-261): HVGK**SSS**RYL | **30** |
| | Trypsin | (residues 1-14): AH-MASMAGPDHLFNLR | 1; 2x: 14 |
| | | (residues 15-42): NHFYLGAYQAAINNSEIPNLSQEDIVER | 23; 2x: 2; 3x: 3 |
| | | (residues 88-95): ESTISSLR | 3 |
| | | (residues 96-110): EWLADPTVGNNAIIR | 2x: 4 |
| | | (residues 188-201): IQEAYLIFQDFSEK | 5 |
| | | (residues 202-212): YPMTSLILNGK | 2 |
| | | (residues 213-235): AVCCMHMGNFEEAETLLLEALNK | 14; 2x: 6 |

**Figure S10. Rhamnosylation site(s) of COPE1 by AvrC.**
PTM analysis of COPE1 and COPE1$^{1-274}$ after incubation with AvrC$^{\Delta 1-51}$. Proteases of AspN, chymotrypsin, or trypsin were used for digestion of the proteins before LC-MS/MS. Peptides identified with modification of +146 Da are listed. Minimum overlapping region for modification is highlighted. S256, S257, and S258 are indicated in red.

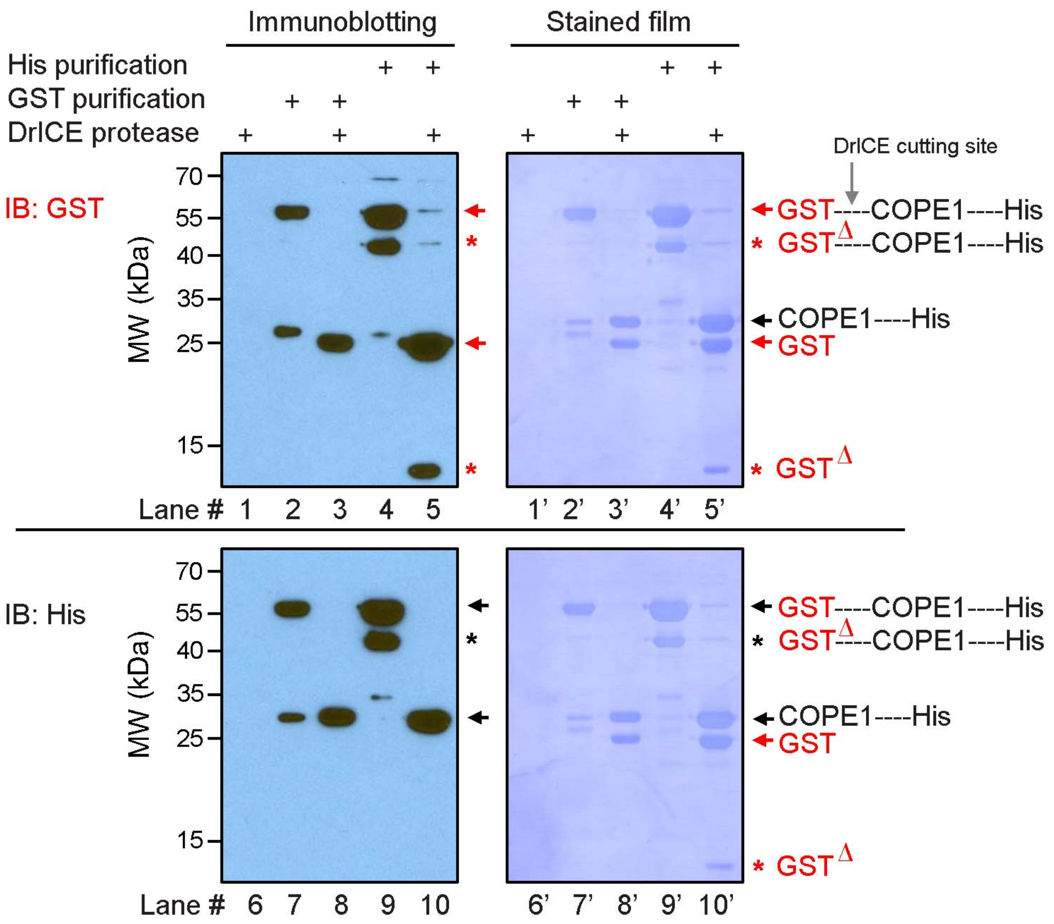


**Figure S11. Purified GST-COPE1-6xHis proteins.**

The proteins were purified using Glutathione agarose against GST tag or Ni-NTA agarose against His tag. Purified proteins were cleaved by DrICE protease to separate GST from COPE1-His. Cleaved products were confirmed by immunoblotting using antibodies against GST and His tag. "*" indicates truncated GST-COPE1-6xHis and GST.

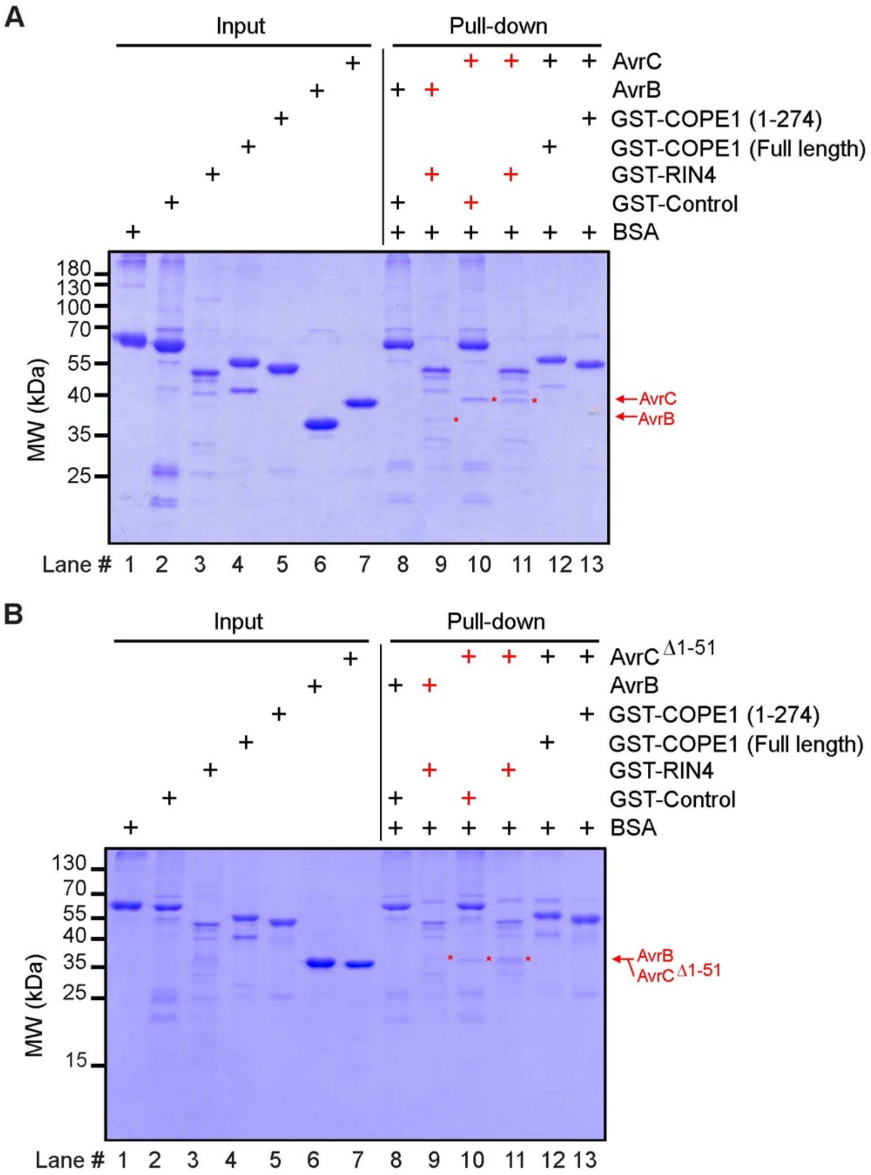


**Figure S12. Interaction test between COPE1 and AvrC in pull-down assays.**
(A, B) Pull-down assay for testing interaction between COPE1 and AvrC or AvrC$^{\Delta 1\text{-}51}$. "+" in red indicates a detected interaction. "*" in red indicates the protein that was pulled down.

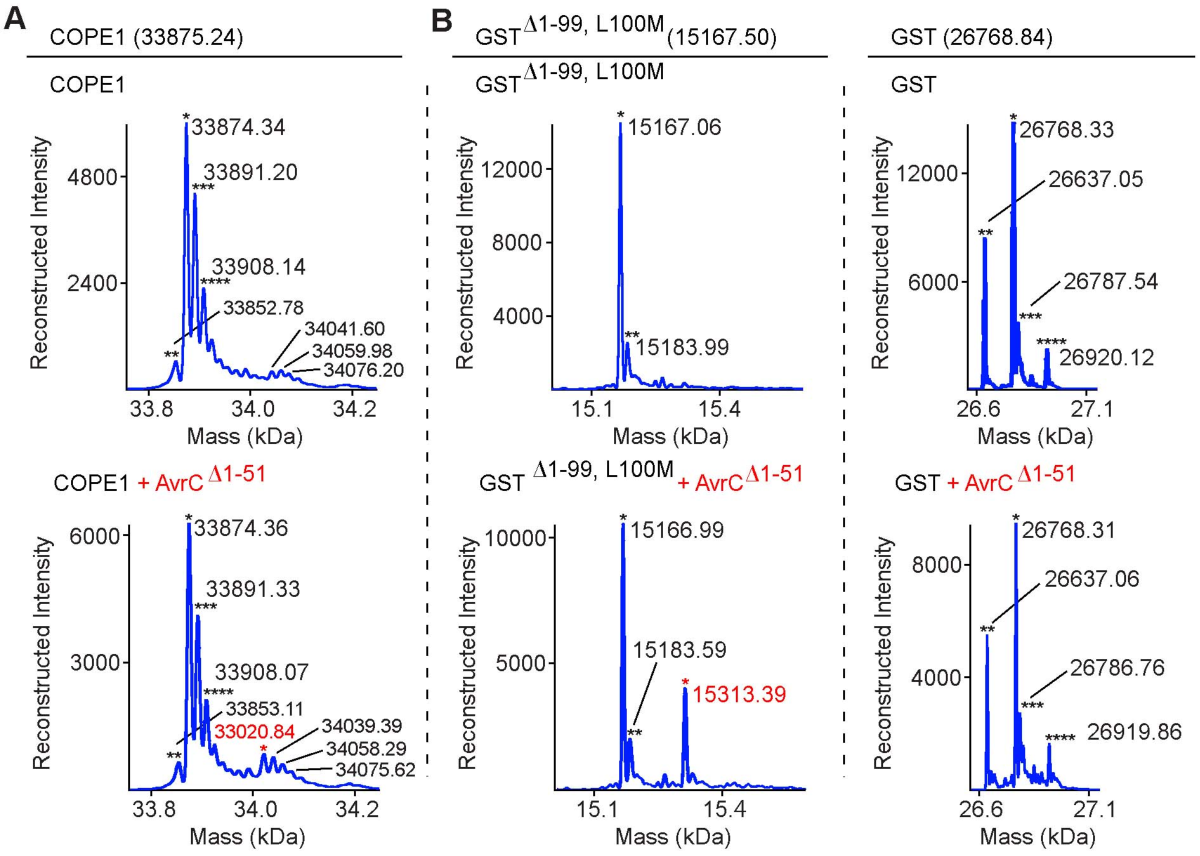


**Figure S13. Rhamnosylation of a truncated GST protein by AvrC$^{\Delta1-51}$.**

(A) Intact mass analysis of COPE1-6xHis after cleavage and incubation with AvrC$^{\Delta1-51}$ in a rhamnosylation assay. "*" symbols in black indicate peaks close to the expected mass. "**", "***", and "****" symbols indicate unknown modification peaks. "*"symbol in red indicates the rhamnosylation peak with a shift of +146 Da.

(B) Intact mass analysis of GST after cleavage and incubation with AvrC$^{\Delta1-51}$ in the rhamnosylation assay in (A). "*" symbols in black indicate peaks close to the expected mass. "**" symbols indicate unknown modification peaks for GST$^{\Delta1-99,\ L100M}$ (left) and peaks likely caused by loss of initiator methionine (-131 Da) for GST (right). "***" and "****" symbols indicate unknown modification peaks for GST (right). "*"symbol in red indicates the rhamnosylation peak with a shift of +146 Da for GST$^{\Delta1-99,\ L100M}$ (left).

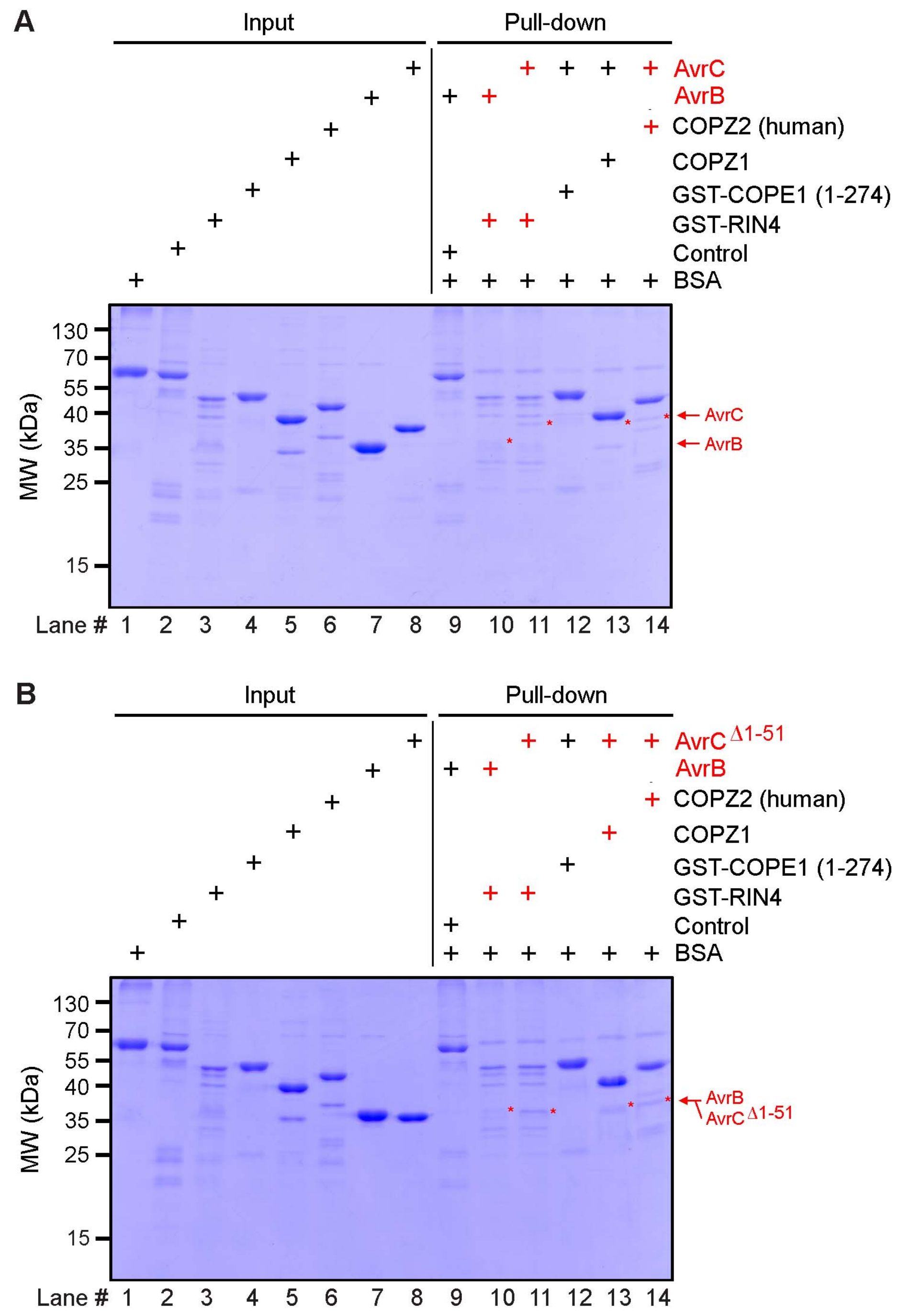


**Figure S14. Interaction between COPZ1 or human COPZ2 and AvrC in pull-down assays.** (A, B) Pull-down assay for testing interaction between COPZ1 or human COPZ2 and AvrC or AvrC$^{\Delta 1\text{-}51}$. “+” in red indicates a detected interaction. “*” in red indicates the protein that was pulled down.

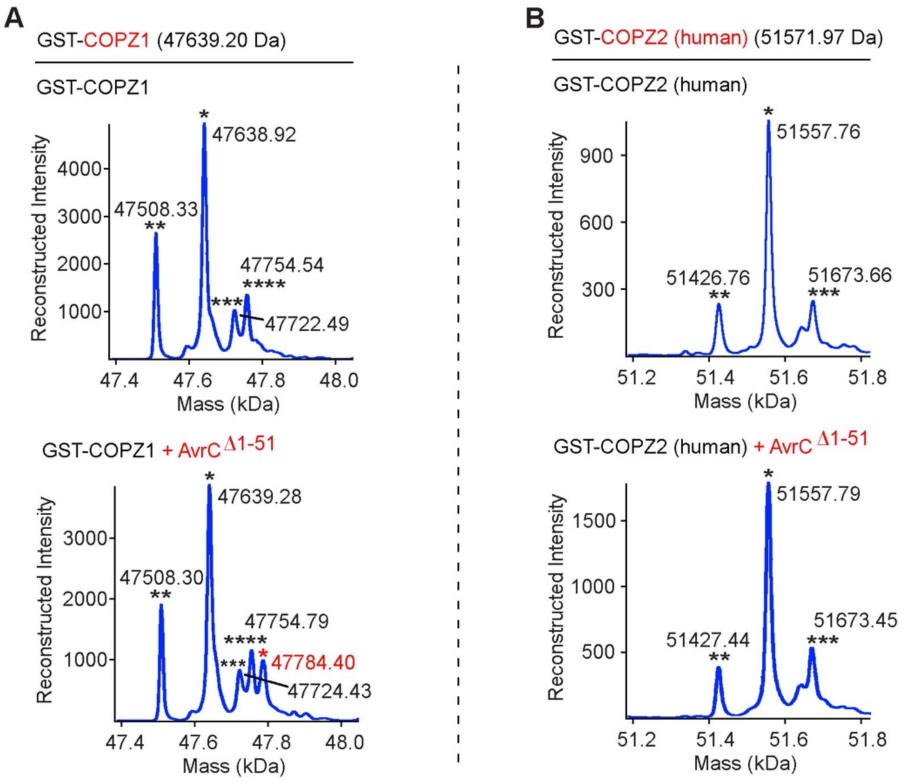


C

| Protein | Protease | Modified peptide (+146 Da) | Hit number |
|---|---|---|---|
| COPZ1 | AspN | (residues 97-107): DAVTLLLRSNV | 2 |
| | Chymotrypsin | (residues 25-43): YSDDWPTNSAQEAFEKSVF | 4 |
| | | (residues 26-43): SDDWPTNSAQEAFEKSVF | 3 |
| | | (residues 134-164): ETDANVIAGKAGINSTDPNAPLSEQTISQAL | 2 |
| | Trypsin | (residues 9-19): NILLLDSEGKR | 2 |
| | | (residues 24-40): YYSDDWPTNSAQEAFEK | 3; 2x: 4 |
| | | (residues 144-168): AGINSTDPNAPLSEQTISQALATAR | 35 |

**Figure S15. Rhamnosylation of COPZ1 by AvrC$^{\Delta 1\text{-}51}$.**

(A, B) Intact mass analysis of GST-COPZ1 and GST-COPZ2 (human) proteins after incubation with AvrC$^{\Delta 1\text{-}51}$ in a rhamnosylation assay.

(C) PTM analysis of COPZ1 after incubation with AvrC$^{\Delta 1\text{-}51}$ in the rhamnosylation assay in (A). Proteases of AspN, chymotrypsin, or trypsin were used for digestion of the proteins before LC-MS/MS. Peptides with modification of +146 Da are listed. Minimum overlapping region for modification is highlighted.

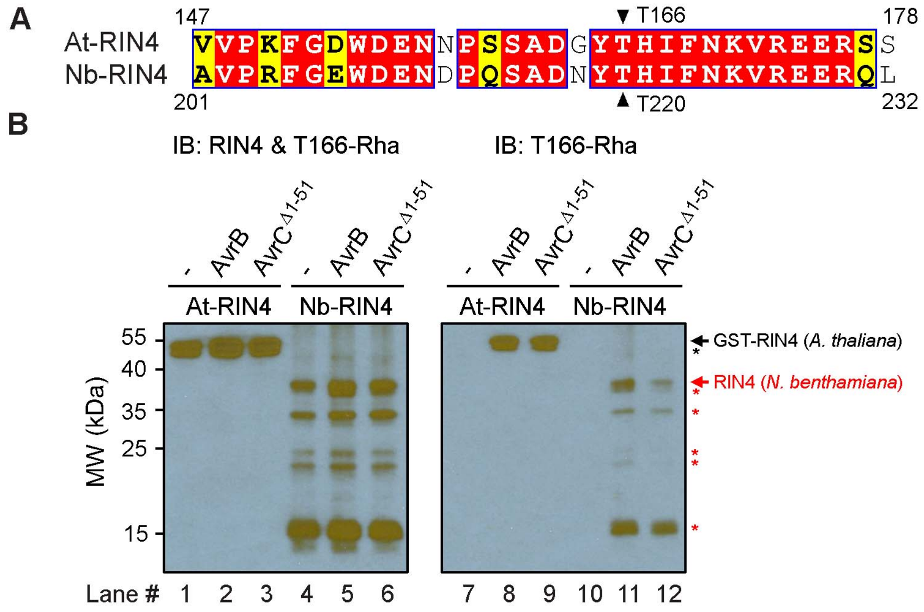


**Figure S16. Rhamnosylation of Nb-RIN4 by AvrB and AvrC$^{\Delta 1\text{-}51}$.**

(A) Alignment of RIN4 from *A. thaliana* and *N. benthamiana*.

(B) *In vitro* rhamnosylation of *N. benthamiana* RIN4 by AvrB and AvrC$^{\Delta 1\text{-}51}$ and immunoblotting using the RIN4$^{T166\text{-}Rha}$ antibodies for detection. "*" indicates likely degraded Nb-RIN4.

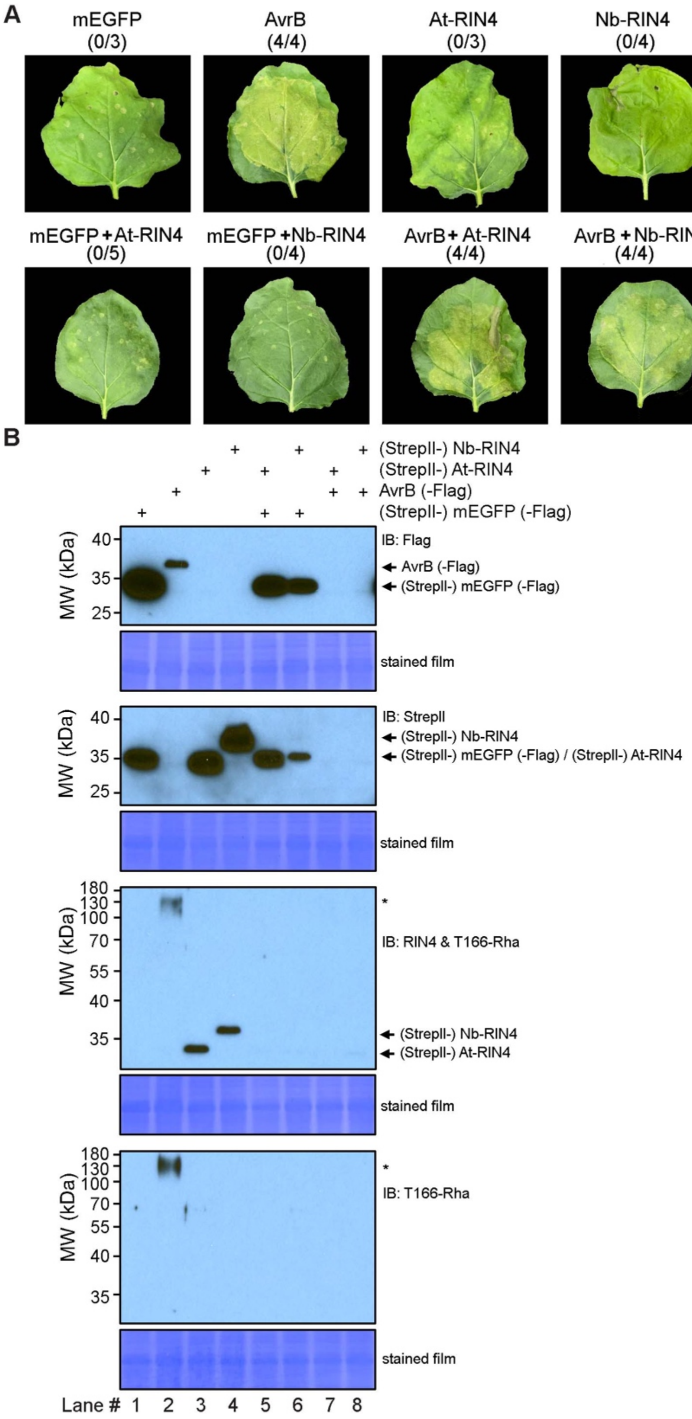


**Figure S17. Tobacco leaf infiltration assay using AvrB.**
(A) *N. benthamiana* leaves were infiltrated by *Agrobacterium* expressing mEGFP (as the vector control), AvrB, *A. thaliana* RIN4 (At-RIN4), and/or *N. benthamiana* RIN4 (Nb-RIN4). Number of leaves showing disease symptoms out of all is indicated in parentheses.
(B) Detection of the expressed proteins using antibodies as indicated. "*" indicates an unknown factor that was likely rhamnosylated by AvrB.

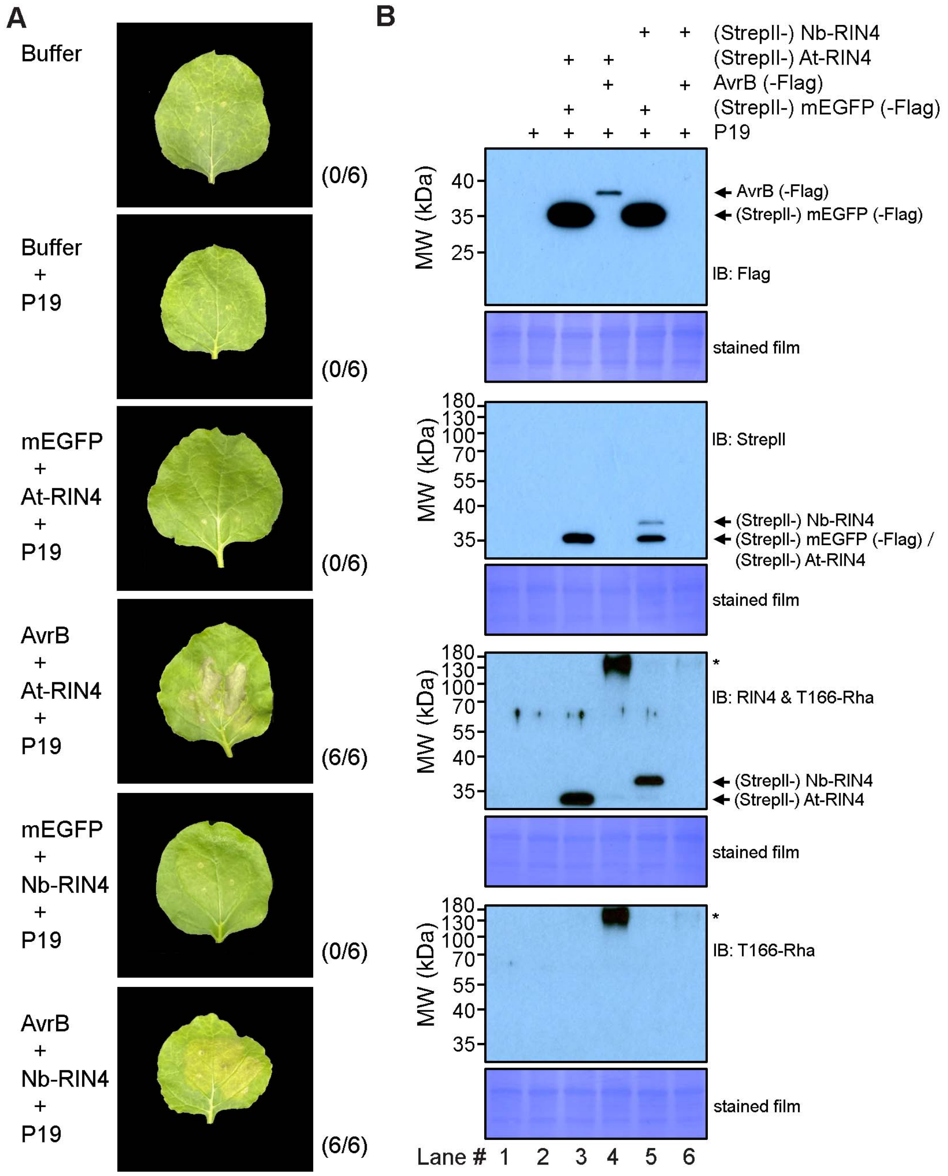


**Figure S18. Tobacco leaf infiltration assay using AvrB and P19 RNA silencing suppressor.**
(A) *N. benthamiana* leaves were infiltrated by *Agrobacterium* expressing mEGFP (as the vector control), AvrB, *A. thaliana* RIN4 (At-RIN4), and/or *N. benthamiana* RIN4 (Nb-RIN4). P19 RNA silencing suppressor was co-expressed to increase protein expression level. Number of leaves showing disease symptoms out of all is indicated in parentheses.
(B) Detection of the expressed proteins using antibodies as indicated. "*" indicates an unknown factor that was likely rhamnosylated by AvrB.

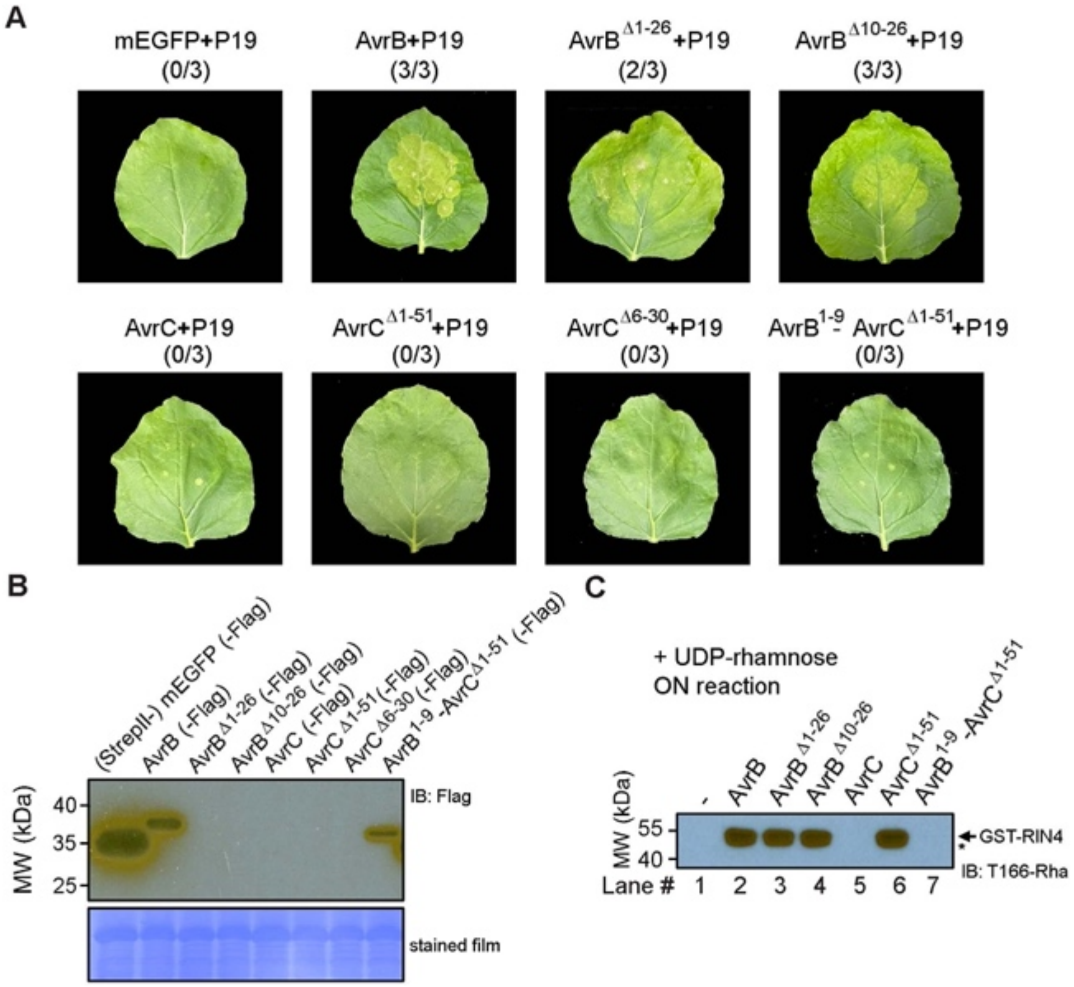


**Figure S19. Tobacco leaf infiltration assay using AvrB and AvrC.**

(A) *N. benthamiana* leaves were infiltrated by *Agrobacterium* expressing mEGFP (as the vector control), AvrB, AvrB$^{\Delta 1-26}$ and AvrB$^{\Delta 10-26}$, AvrC, AvrC$^{\Delta 1-51}$, AvrC$^{\Delta 6-30}$, and AvrB$^{1-9}$-AvrC$^{\Delta 1-51}$. Number of leaves showing disease symptoms out of all is indicated in parentheses.

(B) Detection of the expressed proteins in (A).

(C) *In vitro* rhamnosylation of RIN4 by AvrB and AvrC proteins (purified from BL21(DE3) using UDP-rhamnose as the co-substrate. The reactions were performed overnight (ON). "*" indicates likely degraded RIN4.
